# Downy Mildew effector HaRxL21 interacts with the transcriptional repressor TOPLESS to promote pathogen susceptibility

**DOI:** 10.1101/2020.04.29.066688

**Authors:** Sarah Harvey, Priyanka Kumari, Dmitry Lapin, Thomas Griebel, Richard Hickman, Wenbin Guo, Runxuan Zhang, Jane Parker, Jim Beynon, Katherine Denby, Jens Steinbrenner

**Author notes:** Corresponding author 1: Dr. Jens Steinbrenner, Corresponding author 2: Prof. Katherine Denby. These authors contributed equally to this work (Sarah Harvey, Priyanka Kumari). These authors are joint senior authors on this study (Katherine Denby, Jens Steinbrenner).

## Abstract

*Hyaloperonospora arabidopsidis* (*Hpa*) is an oomycete pathogen causing Arabidopsis downy mildew. Effector proteins secreted from the pathogen into the plant play key roles in promoting infection by suppressing plant immunity and manipulating the host to the pathogen’s advantage. One class of oomycete effectors share a conserved ‘RxLR’ motif critical for their translocation into the host cell. Here we characterize the interaction between an RxLR effector, HaRxL21 (RxL21), and the Arabidopsis transcriptional co-repressor Topless (TPL). We establish that RxL21 and TPL interact via an EAR motif at the C-terminus of the effector, mimicking the host plant mechanism for recruiting TPL to sites of transcriptional repression. We show that this motif, and hence interaction with TPL, is necessary for the virulence function of the effector. Furthermore, we provide evidence that RxL21 uses the interaction with TPL, and its close relative TPL-related 1, to repress plant immunity and enhance host susceptibility to both biotrophic and necrotrophic pathogens.

## Introduction

Plants are constantly under attack by pathogenic microbes. In most cases, the microbe is not able to cause disease on a particular plant species due to pre-formed barriers to infection and/or the ability of the plant to recognize conserved molecular motifs, known as microbe (or pathogen) associated molecular patterns (MAMPs or PAMPs), and activate additional defence responses (Boller & He, 2009; Jones & Dangl, 2006). Signaling pathways activated downstream of MAMP recognition (MAMP or PAMP-triggered immunity; PTI) by pattern recognition receptors (PRRs) result in production of reactive oxygen species, hormone biosynthesis, callose deposition in the cell wall and large-scale transcriptional reprogramming within the plant (Kunze et al., 2004; Navarro et al., 2004). Many pathogens use effector proteins to suppress or evade these host immune responses and/or adapt host physiology to aid infection (Toruño, Stergiopoulos, & Coaker, 2016). For example, effectors may target PTI signalling (de Jonge et al., 2010; Feng et al., 2012; P. He et al., 2006; Shang et al., 2006) or manipulate stomatal opening (Gimenez-Ibanez et al., 2014). Manipulation of the host plant by the pathogen also occurs through alteration of host transcription; it is well documented that transcription activator-like effector (TALe) proteins from the *Xanthomonas* genus of plant pathogenic bacteria bind and activate host promoters (Römer et al., 2010). Large-scale changes in host transcription brought about by effectors have also been shown during infection by *Pseudomonas syringae* pv. *tomato* DC3000 (*Pst*) (Lewis et al., 2015; Thilmony, Underwood, & He, 2006).

Alignment of known effector proteins from plant pathogenic oomycetes including *Hyaloperonospora arabidopsidis* (*Hpa*) revealed a consensus sequence RxLR (arginine, any amino acid, leucine, arginine) downstream from a signal peptide (Rehmany et al., 2005), facilitating identification and subsequent characterization of candidate effector proteins from the *Hpa* genome (Baxter et al., 2010; Fabro et al., 2011). Potential Arabidopsis protein targets of *Hpa* effectors have been identified using yeast-2-hybrid (Y2H) with 122 different Arabidopsis proteins targeted by 53 *Hpa* effectors (Mukhtar et al., 2011; Weßling et al., 2014). One effector from *Hpa*, HaRxL21 (RxL21) was found to interact with the Arabidopsis transcriptional corepressor TOPLESS (TPL) (Mukhtar et al., 2011).

In Arabidopsis, the TPL family consists of five members, TPL and four TPL-related (TPR) proteins. TPR1 is the most similar to TPL sharing 95% similarity at the amino acid level (Kagale & Rozwadowski, 2011; Long, Ohno, Smith, & Meyerowitz, 2006; Zhu et al., 2010). TPL and TPRs have been shown by Y2H to interact with several regulators of hormone pathways known to be involved in plant defence against pathogens (Arabidopsis Interactome Mapping Consortium, 2011), abiotic stress and in development (Causier, Ashworth, Guo, & Davies, 2012a). Moreover, involvement of TPL/TPRs in regulation of resistance protein signalling (Van Den Burg & Takken, 2010; Zhu et al., 2010) and jasmonate signalling (Pauwels et al., 2010) has been shown. We speculate that TPL is a key target for manipulation by pathogens.

The TPL family is highly conserved in land plants (Causier, Lloyd, Stevens, & Davies, 2012b; Hao et al., 2014) and shows structural similarity to the *Drosophila* protein Groucho, in addition to human transducin-like Enhancer of split and transducin (beta)- like 1 proteins (G. Chen & Courey, 2000; Martin-Arevalillo et al., 2017; Oberoi et al., 2011). TPL family members link transcription factors (TFs) to chromatin remodelling complexes; the corepressors interact with TFs and recruit chromatin remodelling factors such as histone deacetylases (Kagale & Rozwadowski, 2011; Wang, Kim, & Somers, 2013; Zhu et al., 2010). For example, the Arabidopsis regulators of root hair development GIR1 and GIR2 have been shown to promote histone hypoacetylation via their interaction with TPL (R. Wu & Citovsky, 2017). The current understanding is, therefore, that TPL/TPR recruitment to a gene results in reversible transcriptional repression via condensed DNA structure that is less accessible to transcription initiation complexes.

At the very N-terminus TPL harbours a LIS1 homology domain (LisH) followed by a C-terminal to LisH (CTLH) domain. The latter is needed for the interaction with proteins containing an <u>E</u>thylene-responsive element Binding Factor-associated <u>A</u>mphiphilic <u>R</u>epression (EAR) motif (Szemenyei, Hannon, & Long, 2008; Zeng et al., 2006). The EAR motif was first identified as the conserved sequence ^L^/_F_DLN^L^/_F_ (x)P in class II Ethylene Response Factor genes which function as transcriptional repressors (Ohta, Matsui, Hiratsu, Shinshi, & Ohme-Takagi, 2001). EAR-mediated protein-protein interaction with TPL is often required for transcriptional repression. For example, the transcriptional regulator IAA12 requires interaction between its EAR motif and the CTLH domain of TPL for its repressive activity in low auxin conditions (Szemenyei et al., 2008; Zeng et al., 2006). Repression of jasmonate signalling relies on interaction between TPL and the EAR motif-containing Novel Interactor of JAZ (NINJA) (Pauwels et al., 2010; Pérez & Goossens, 2013). Interaction with TPL and transcriptional repressor activity of NINJA were abolished when Leu residues in the EAR motif were mutated to Ala (Pauwels et al., 2010).

EAR motifs have now been identified in many plant proteins involved in development, stress and defence (C.-J. Dong & Liu, 2010; Espinosa-Ruiz et al., 2017; Krogan, Hogan, & Long, 2012). So far, a few pathogen effectors mimicking this EAR motif have been described although little is known about subsequent corepressor recruitment. For example, the *Xanthomonas campestris* type III effector XopD has been found to contain two tandemly repeated EAR motifs downstream of a DNA binding domain which are required for XopD-dependent virulence in tomato (Canonne et al., 2011; J.-G. Kim, Taylor, & Mudgett, 2011). Effectors from the XopD superfamily that contain conserved EAR motifs have been found in *Xanthomonas, Acidovorax* and *Pseudomonas* species (J.-G. Kim et al., 2011). In addition, a conserved EAR motif is required for virulence in the *Ralstonia solanacearum* effector PopP2 although again no interaction with known Arabidopsis corepressors has been identified so far (Segonzac et al., 2017). There are however, examples of pathogen effectors interacting with TPL family members including interaction of the *Melampsora larici-populina* effector MLP124017 with TPR4 (Petre et al., 2015).

The aim of our work was to determine how RxL21 is manipulating Arabidopsis to promote infection by *Hpa*. We demonstrate that RxL21 interacts *in planta* with the Arabidopsis corepressors TPL and TPR1 via an EAR motif and that this interaction is essential for RxL21 virulence activity against both Hpa and a necrotrophic plant pathogen. We find there is co-occurrence of RxL21 and TPR1 binding sites on promoter regions of a set of TPR1-repressed defence-related genes, suggesting that RxL21 virulence function involves perturbation of TPL/TPR1 transcriptional repression during mobilization of host immunity.

## Results

### RxL21 is conserved across multiple *Hpa* isolates and contains a C-terminal EAR motif

RxL21 is a 45 kDa effector protein identified from the genome of *Hpa* (Fabro et al., 2011). It contains a ‘RLLR-DEER’ motif at the N-terminus and an EAR motif (amino acid sequence LMLTL) at the C-terminus. Alleles of *RxL21* have been found in *Hpa* isolates Cala2, Emco5, Emoy2, Emwa1, Hind2, Maks9, Noks1 and Waco9 ((Asai et al., 2018) and BioProject PRJNA298674). Alignment of the amino acid sequences of RxL21 alleles was performed using T-COFFEE (Di Tommaso et al., 2011) (Fig S1). The signal peptide cleavage site was predicted to be between position 16 and 17 (SignalP-5.0). The RxLR-DEER motif is conserved across all alleles and with the exception of Noks1 (truncated due to Serine 197 changed to a Stop codon) the EAR motif at the C-terminus is also conserved between all aligned alleles of RxL21. *RxL21* is expressed in conidiospores of virulent (Waco9) and avirulent (Emoy2) *Hpa* isolates during Col-0 infection, as well as in Waco9 1, 3 and 5 days post inoculation (Asai et al., 2014) consistent with it playing a role in virulence.

### RxL21 expression *in planta* causes enhanced susceptibility to both biotrophic and necrotrophic pathogens

Previously, during screening of multiple candidate effectors from *Hpa*, constitutive expression of RxL21 (from *Hpa* isolate Emoy2) in Arabidopsis has been shown to enhance growth of *Hpa* isolate Noco2 and a *Pst* isolate impaired in the suppression of early immune responses (*Pst* DC300 ΔavrPto/ΔavrPtoB) (Fabro et al., 2011). In addition, RxL21 increased bacterial growth when delivered into Arabidopsis via the type-three secretion system of *Pst* DC3000-LUX (Fabro et al., 2011). To assess whether the pathogen susceptibility boost provided by RxL21 expression *in planta* extends to both biotrophic and necrotrophic pathogens, Arabidopsis plants expressing RxL21 under the control of a 35S promoter (Fabro et al., 2011) were screened for susceptibility to *Hpa* isolates Noks1 and Maks9, and the necrotrophic pathogen *Botrytis cinerea*. RxL21 lines were compared to both Col-0 wild type and lines expressing *35S::GUS* (Col-0 GUS) which had been transformed and selected alongside the effector lines. Presence of the effector was found to confer enhanced susceptibility to both obligate biotroph *Hpa* isolates in two independent transgenic lines (RxL21a, b) compared to both Col-0 and Col-0 GUS, measured by total sporangiophores per seedling at 4 days post inoculation (dpi) (Fig 1A). RxL21 expression also resulted in increased lesion size caused by *B*. *cinerea* infection compared to controls (Fig 1B). Hence, it appears that the RxL21 effector is targeting a mechanism essential for a full defence response against both biotrophic and necrotrophic pathogens.

**Figure 1.**
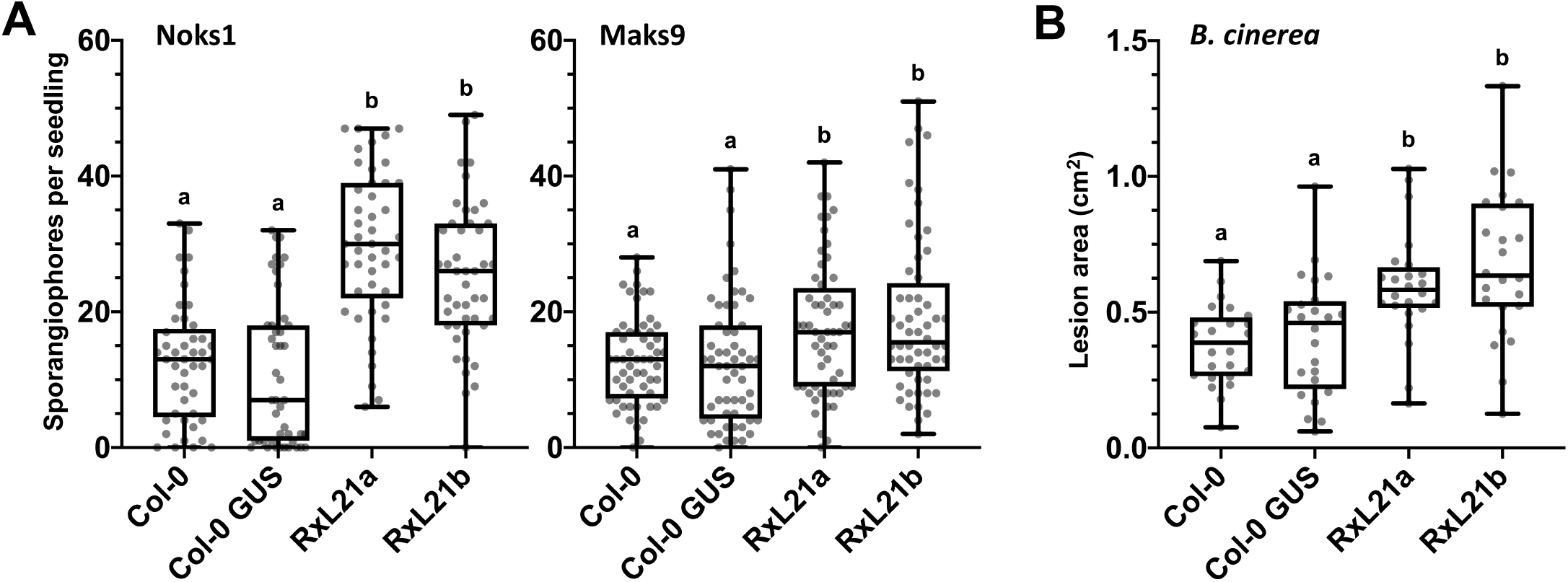
RxL21 expression *in planta* causes enhanced susceptibility to biotrophic and necrotrophic pathogens. (A) Transgenic Arabidopsis expressing RxL21 under a 35S promoter (RxL21a/b) were challenged with *Hpa* isolates Noks1 and Maks9 and sporangiophores counted at 4 dpi. Col-0 WT and 35S::GUS (Col-0 background) were used as controls. (Noks1 n=45, Maks9 n=55). (B) RxL21a/b were challenged with *B*. *cinerea* and lesion area measured 72 h post infection (n=24). Box plots show the median, upper and lower quartiles, whiskers show the upper and lower extremes of the data. Letters indicate significant difference using a Kruskal Wallis test with p < 0.05. Experiments were repeated with similar results.

### RxL21 interacts with the CTLH domain of TPL via its EAR motif

It has been previously reported that RxL21 interacts with TPL using the matrix yeast two hybrid technique (Y2H) (Mukhtar et al., 2011). To confirm that this interaction could occur *in planta*, we first determined that RxL21 and TPL co-localise to the nucleus when transiently expressed in *N*. *benthamiana* leaves (Fig 2A). We next investigated the specific interaction motifs of both partners, which have not been previously reported. Multiple truncated forms of RxL21 with or without the EAR motif and/or RxLR motif were cloned and used in Y2H analysis with TPL. Two EAR-mutation constructs (RxL21ΔEAR1; Δ402-409 and RxL21ΔEAR2; Δ360-409) were used to determine whether the amino acids flanking the EAR motif are necessary for interaction with TPL (Fig 2B). As expected, full-length RxL21 showed a positive interaction with TPL via activation of histidine (*GAL1::HIS3*) and adenine (*GAL2::ADE2*) reporter genes. The screen was performed in both directions with RxL21 and TPL fused to both the activation domain (AD) or the DNA binding domain (DB). Selective plates containing 3AT (a competitive inhibitor of the HIS3 gene product) were used as an additional control for increased stringency. Binding affinity appeared to be influenced by the direction of cloning with activation of the adenine reporter gene (and high stringency activation of the histidine reporter) only detected when TPL was fused to the DB domain. Deletion of either the initial N terminal sequence (RxL21ΔN) or the RxLR-DEER motif (RxL21ΔRxLR) did not prevent interaction with TPL. However, deletion of the EAR motif abolished the interaction with TPL (Fig 2B), demonstrating that the EAR motif is essential for this direct protein-protein interaction in yeast. Deletion of the CTLH domain of TPL abolished the interaction with RxL21 (Fig 2C). Hence, these data show that both the EAR motif of RxL21 and the CTLH domain of TPL are necessary for direct protein-protein interaction. The mechanism of interaction between the effector and its host target (TPL) is therefore mimicking the mechanism by which plant proteins (such as NINJA (Pauwels et al., 2010)) interact with TPL and recruit it to a complex for transcriptional repression.

**Figure 2.**
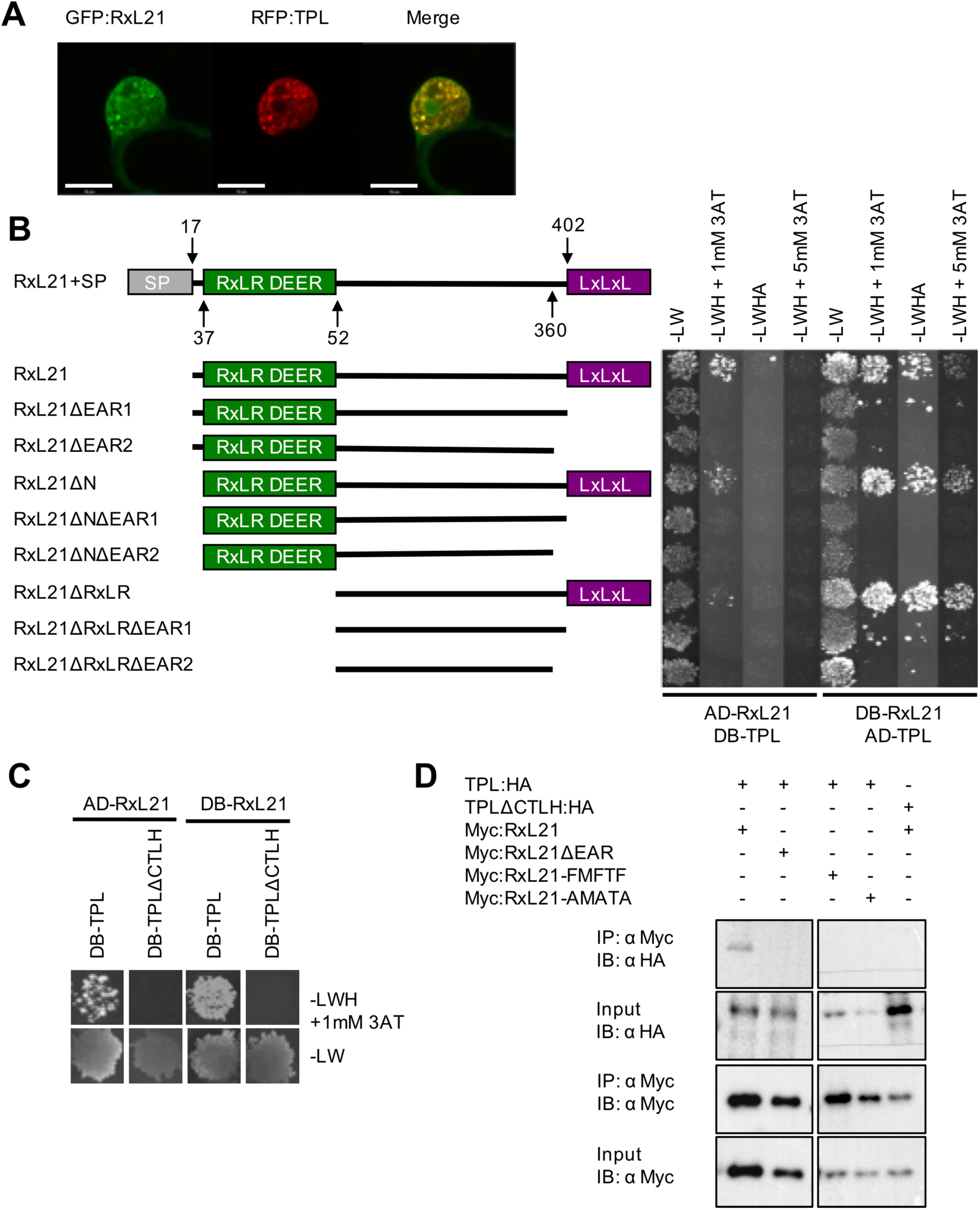
The EAR motif of RxL21 interacts with the CTLH domain of TPL. (A) RxL21 co-localizes with TPL in the nucleus. GFP:RxL21 and RFP:TPL were expressed transiently in *N*. *benthamiana*. Scale bars are 10 μm. (B) TPL Interacts with the EAR domain of RxL21 in yeast. RxL21 was cloned without the signal peptide and lacking the EAR motif (RxL21ΔEAR1; Δ402-409 and HaRxL21ΔEAR2; Δ360-409), without the N-terminus (RxL21ΔN; Δ1-36) and RxLR-DEER motif (RxL21ΔRxLR; Δ1-51) and combinations of the above. Amino acid locations of deletion constructs are indicated. Deleting the EAR motif at the C-terminus of RxL21 abolishes interaction with TPL by Y2H. Successful mating is indicated by growth on media lacking Leucine and Tryptophan (-LW). Growth on media additionally lacking Histidine (-LWH) indicates *GAL1::HIS3* reporter gene activation and protein-protein interaction. 3AT is used to increase stringency in a concentration dependent manner. Growth on media additionally lacking Adenine (-LWHA) indicates activation of the *GAL2::ADE2* reporter gene. AD; activation domain. DB; DNA binding domain. (C) Deleting the CTLH domain of TPL (TPLΔCTLH; Δ25-91) abolishes interaction with RxL21 in yeast. Interaction (indicated by growth on –LWH +1mM 3AT) was observed between RxL21 and TPL but not between RxL21 and TPLΔCTLH. All Y2H was repeated at least twice with similar results. (D) TPL interacts with RxL21 and the interaction requires the EAR motif *in planta*. TPL:HA, TPLΔCTLH:HA together with RxL21 and RxL21 EAR motif mutants RxL21ΔEAR, RxL21-FMFTF and RxL21-AMATA (under control of an estradiol-inducible promoter and N-terminal Myc tag) were transiently expressed in *N*. *benthamiana* leaves. RxL21 expression was induced by 30μM β-estradiol 24 hr prior to harvesting. c-myc beads were used for immunoprecipitation (IP). HA antibody was used for TPL and TPLΔCTLH immunoblots (IB) and c-myc antibody was used to detect RxL21, RxL21ΔEAR and the respective EAR motif mutants.

### Leucine residues in the EAR motif are critical for interaction with TPL

In addition to deletion of the EAR motif (RxL21ΔEAR), site-directed mutagenesis of the EAR motif was performed to assess the importance of individual amino acid residues. The three Leucine (Leu/L) residues (Leu389, Leu391, Leu393) in the EAR motif of RxL21 were mutated individually, in pairs or all together to Phenylalanine (Phe/F), Alanine (Ala/A) and Isoleucine (Ile/I). Substitution to Phe was chosen because previously Leu to Phe mutation of Leu354, Leu356 and Leu358 in the Arabidopsis repressor SUPERMAN prevented corepressor activity (Hiratsu, Mitsuda, Matsui, & Ohme-Takagi, 2004). In addition, the three Leu residues in RxL21 were mutated to Ala (as it lacks functional side chains) (Morrison & Weiss, 2001) and Ile because it is the most similar amino acid to Leu (Livingstone & Barton, 1993).

Mutagenesis of all three Leu residues to Phe, Ala or Ile was found to abolish interaction of RxL21 with TPL by Y2H (Fig S2). Mutation of the ‘x’ residues within the EAR motif between the Leu residues (Methionine390 and Threonine392) had no effect on the interaction. Mutation of the first Leu residue (Leu389) or any pair of Leu residues in the EAR motif was sufficient to abolish RxL21-TPL interaction (Fig S2A). This analysis was repeated using TPL from *Nicotiana benthamiana* (*Nb*TPL). RxL21 interacted with *Nb*TPL and the same residues in the RxL21 EAR motif were necessary for interaction with *Nb*TPL as for interaction with *At*TPL (Fig S2B). These data are consistent with previous observations that mutation of any two Leucine residues within the DLELRL hexapeptide of SUPERMAN is sufficient to abolish repression of transcription (Hiratsu et al., 2004).

### RxL21 and TPL interact *in planta*

Having demonstrated the importance of the EAR motif and CTLH domain of RxL21 and TPL respectively for interaction in yeast, we next verified that this protein interaction occurs *in planta*. For bimolecular fluorescence complementation (Split YFP) RxL21 was cloned into a vector with an N-terminal fused YFP fragment to minimise steric hindrance to the C-terminal EAR motif. *Agrobacterium tumefaciens* harbouring the N and C fragments of E-YFP fused to RxL21 and TPL were infiltrated into *N*. *benthamiana* and imaged using confocal microscopy. YFP fluorescence was detected in the nucleus, but no fluorescence was observed when RxL21ΔEAR was used instead of the full-length effector (Fig S3).

The interaction between RxL21 and TPL was also confirmed by co-immunoprcipitation (Co-IP) using *A*. *tumefaciens* mediated transient expression in *N*. *benthamiana*. A Myc tag was fused to the N terminal of RxL21 and the construct co-expressed with C-terminal HA-tagged TPL. Immunoprecipitation of RxL21 using an anti-Myc antibody resulted in Co-IP of TPL (Fig 2D) confirming direct protein-protein interaction occurs *in planta*. Furthermore, Myc-tagged RxL21ΔEAR was unable to pull down TPL, and TPLΔCTLH was not immunoprecipitated by Myc-tagged RxL21. This demonstrated that, as in Y2H, the EAR motif and CTLH domain are required for RxL21-TPL interaction. TPL was also not immunoprecipitated when using variants of RxL21 in which the Leu residues in the EAR motif were mutated to Phe or Ala (RxL21-FMFTF or RxL21-AMATA), again confirming our observations in yeast that the Leu residues in the EAR motif of RxL21 are necessary for interaction with TPL.

### Additional *Hpa* effectors containing EAR motifs do not interact with TPL

Having established the importance of the EAR motif for interaction with TPL, we investigated whether other *Hpa* effectors contained this motif and could interact with TPL. In a previous large-scale study, no other *Hpa* effectors were found to interact with TPL (Mukhtar et al., 2011). The amino acid sequence of 134 RxLR (RxL), 491 RxL-like (RxLL) and 20 Crinkler (CRN) candidate effector proteins in the *Hpa* genome (Asai et al., 2018; Baxter et al., 2010) were searched for the presence of the LxLxL motif. The predicted signal peptide cleavage site in the effectors was then identified using SignalP (Petersen, Brunak, Heijne, & Nielsen, 2011) and any effectors containing an LxLxL motif within the predicted signal peptide were subsequently excluded from further analysis.

The LxLxL motif was found to be present in 16 RxLs and 35 RxLLs (Table S1). Of these, 2 RxLs and 4 RxLLs were found to contain multiple LxLxL motifs. The position of the LxLxL motif was noted, since previous reports of the EAR motif in functionally characterized transcriptional repressors is often at the N or C terminus (Pauwels et al., 2010; Shyu et al., 2012; Tiwari, Hagen, & Guilfoyle, 2004). The LxLxL motif of 5 RxLs (including RxL21) and 7 RxLLs was found to be within 35 amino acids of the C-terminus of the protein. No RxLs or RxLLs were found to have a LxLxL motif at the N terminus, except those which were predicted to be within the signal peptide. We compared LxLxL-containing effectors against available subcellular localization data (Caillaud et al., 2011) and found, in addition to RxL21, HaRxL48, HaRxL55, HaRxLL100 and HaRxLL470 were nuclear localised in *Nicotiana benthamiana* (Caillaud et al., 2011), suggesting a possible functional role for the EAR motif in these effectors in manipulating host transcription. Ten of the LxLxL-containing *Hpa* effectors (including three of the four nuclear-localised ones) were tested using Y2H. None of the effectors showed an interaction with TPL (Fig S4) indicating that the presence of an EAR motif is not sufficient to mediate an interaction with TPL and that RxL21 may have a unique function.

### TPL family members function in defence against multiple pathogens

It has previously been shown that TPL, TPR1 and TPR4 function in defence against *Pst*, with redundancy observed between TPL and TPR1 (Zhu et al., 2010). We sprayed *tpr1*-*tpl*-*tpr4* triple knockout plants, as well as Col-0 and Col-0 35S::GUS controls with *Hpa* isolate Noks1 and counted sporangiophores per seedling at 4 dpi. The *tpr1*-*tpl*-*tpr4* plants were found to be significantly more susceptible (Fig 3A) than both wild type (Col-0) and Col-0 GUS control plants. In addition, *tpr1*-*tpl*-*tpr4* plants were drop inoculated with *B*. *cinerea* and showed significantly increased lesion area (Fig 3B) and visual symptoms of sporulation (Fig 3C). These data suggest that transcriptional regulation by TPL and family members is a crucial part of immune signalling against multiple pathogens with different lifestyles.

**Figure 3.**
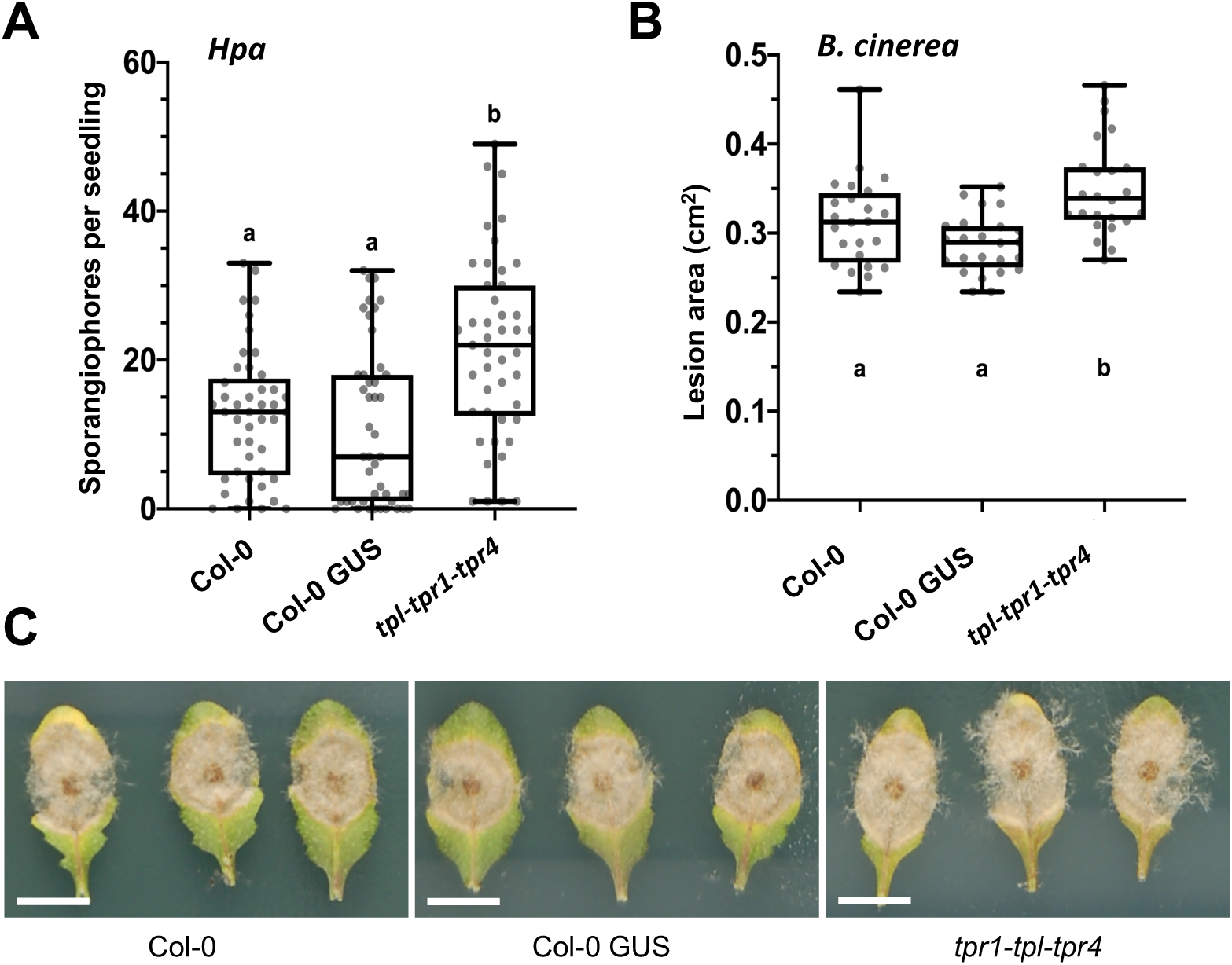
TPL family members function in immunity against a biotrophic and a necrotrophic pathogen. (A) *tpl-tpr1-tpr4* plants showed enhanced susceptibility to *Hpa* isolate Noks1 measured by sporangiophore counts per seedling at 4 dpi compared to Col-0 WT and 35s::GUS controls. Letters indicate significance using a Kruskal Wallis test, n=45, p < 0.05. (B) Enhanced susceptibility of *tpl-tpr1-tpr4 to B*. *cinerea* infection was also observed as determined by lesion area at 48 hpi (n=24, letters indicate significance using a one-way ANOVA and a Tukey test, p < 0.05) and (C) visual inspection of sporulation. Here we show representative leaf images at 96h post infection. Scale bar is 1 cm.

### RxL21 also interacts with TPR1

The library used in the original Y2H screen identifying the interaction between RxL21 and TPL did not include all members of the TPL corepressor family in Arabidopsis (Arabidopsis Interactome Mapping Consortium, 2011). TPL/TPR interactions can exhibit both redundancy (eg. all family members interact with multiple IAAs) and specificity (eg. several ERF TFs interact with 1 member or a subset of the TPL/TPR family (Causier, Ashworth, Guo, & Davies, 2012a)). We tested interaction between all four TPRs and RxL21 using Y2H. In addition to TPL, RxL21 was found to interact with TPR1 (Fig S5A), the most closely related family member to TPL with 95% similarity at the amino acid level (Zhu et al., 2010). As with TPL, this interaction was abolished when the EAR motif of RxL21 was deleted. The RxL21-TPR1 interaction was also confirmed by Co-IP (Fig S5B) using transient expression in *N*. *benthamiana*.

### The EAR domain is necessary for RxL21 virulence function

To determine whether interaction with TPL is required for the ability of RxL21 to enhance susceptibility of Arabidopsis to pathogens, homozygous transgenic plants were generated expressing either RxL21 or RxL21 with a truncation to remove the EAR motif. In these lines, RxL21 was fused to a N-terminal HA tag and expressed in a Col-4 background under the control of the 35S promoter (HA:21#1/2 or HA:21ΔEAR#1/2). Expression of the transgenes was verified by qPCR and western blot (Fig S6). We noted that protein expression levels in all transgenic lines were very low perhaps due to a detrimental effect of effector accumulation. However, given the effector mRNA we challenged these lines, compared to Col-4 WT and HA:GFP (35S promoter, Col-4 background) control plants, to *Hpa* infection. The HA:21 lines exhibited enhanced susceptibility whereas the HA:21ΔEAR lines did not differ in susceptibility from WT and GFP controls (Figure 4A). Similarly, compared to HA:GFP controls, both HA:21 lines showed significantly enhanced lesion area at 72 hpi after inoculation with *B*. *cinerea* whereas HA:21ΔEAR lines did not (Fig 4B). Enhanced sporulation was also clearly seen on leaves from the HA:21#1 line compared to Col-4 WT, HA:GFP controls and HA:21ΔEAR#1 line (Fig S7). These data demonstrate that the EAR motif of RxL21 (and hence most likely interaction with TPL/TPR1) is necessary for enhanced susceptibility to biotrophic and necrotrophic pathogens.

**Figure 4.**
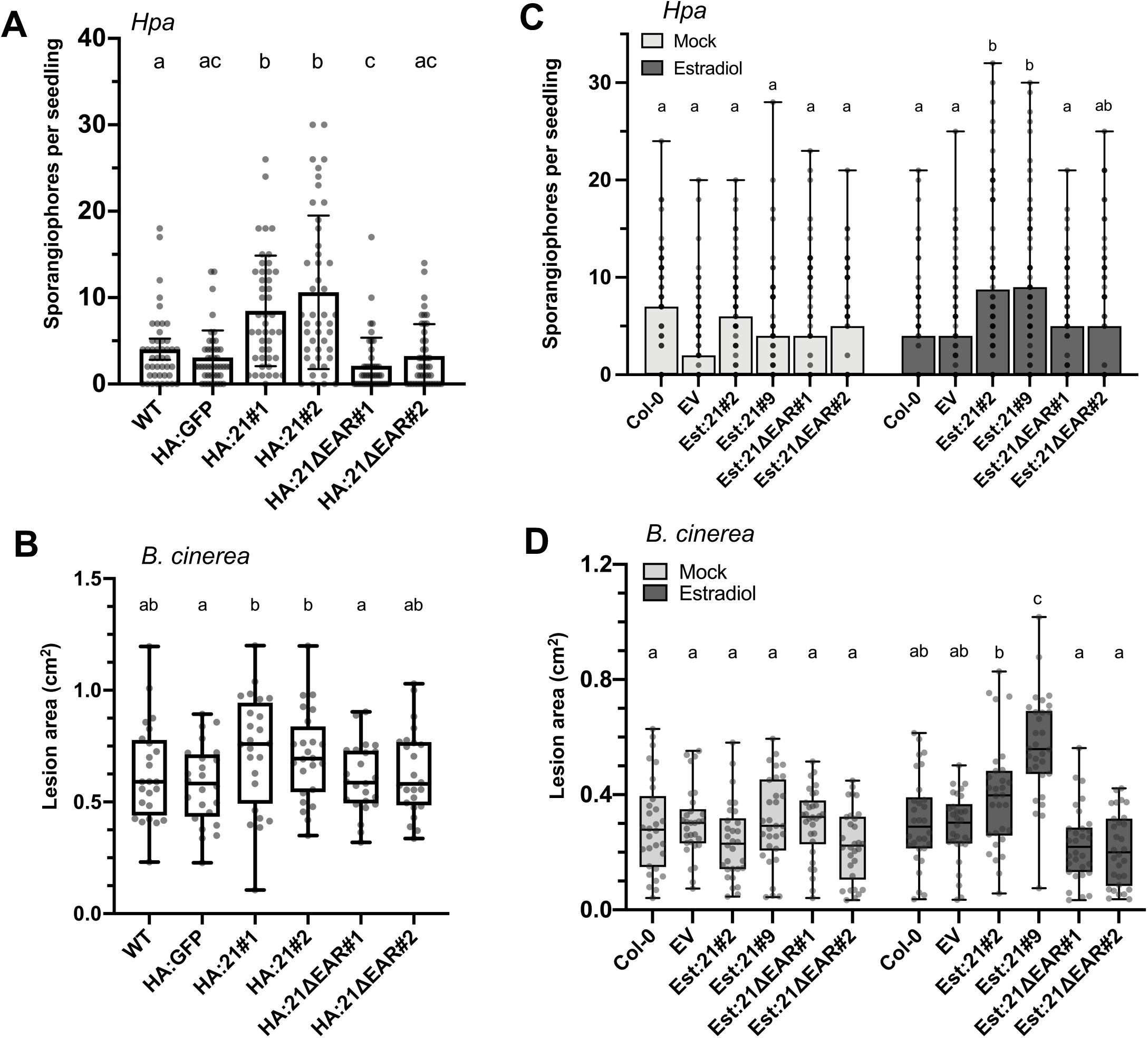
The EAR motif is necessary for RxL21 virulence function *in planta*. Transgenic lines expressing RxL21 under control of a 35S promoter and N-terminal HA tag (HA:21#1/2 and HA:21ΔEAR#1/2) were compared to Col-4 WT and HA:GFP (Col-4 background) controls during infection with (A) *Hpa* isolate Noks1 (sporangiophore counts per seedling at 4 dpi). Letters indicate significance using a Kruskal Wallis test, p < 0.05, n=45 (B) *B*. *cinerea*. Lesion area was measured at 72 hpi (letters indicate significance determined by a one way ANOVA, p < 0.05, n=24). Error bars display 95% CE. (C) Estradiol inducible lines expressing RxL21 and RxL21ΔEAR fused to an N-terminal Myc tag (Est:21#2/9 and Est:21ΔEAR#1/2) were infected with *Hpa* 18 hours after induction with 30 μM β-estradiol or mock treatment (n=100 per treatment). (D) The same lines with estradiol or mock treatment were subsequently infected with *B*. *cinerea*. Lesion area was measured at 72 hpi (n=30). Letters indicate significant difference between treatments using a 2-way ANOVA and Bonferroni’s multiple comparison test. P < 0.05. Whiskers show data range. Experiments were repeated with similar results.

To verify that the enhanced susceptibility of HA:21 lines was directly due to effector expression rather than physiological or developmental differences between transgenic and control plants due to continual overexpression of the effector, we generated lines expressing RxL21 and RxL21ΔEAR under an estradiol-inducible promoter and with an N-terminal fused Myc tag (Est:21#2/6/9 and Est:21ΔEAR#1/2/3). Expression of RxL21 after estradiol induction was verified by qPCR and protein accumulation (before and after induction) was assessed by western blot (Fig S8). After estradiol induction, Est:21 lines showed enhanced susceptibility to *Hpa* infection (as measured by sporangiophore counts; Fig 4C) and *B*. *cinerea* infection (as measured by lesion area; Fig 4D). In contrast, Est:21ΔEAR lines did not show enhanced susceptibility to either pathogen, confirming that the EAR motif is necessary for the virulence function of the effector and that expression of the effector shortly before infection is sufficient to enhance host susceptibility.

### RxL21 interaction with TPL affects host transcription

Both the nuclear localisation and protein-protein interaction with TPL within the nucleus suggest that RxL21 plays a role in manipulation of host transcription. We observed that when GFP::RxL21 is transiently expressed in *N*. *benthamiana*, treating with DAPI before imaging results in co-localisation of GFP and DAPI (Fig S9). DAPI has been shown to accumulate with high levels of heterochromatin (Linhoff, Garg, & Mandel, 2015) and therefore tightly bound chromatin in a repressed state. This observation is consistent with our observations that RxL21 is able to interact with TPL from *N*. *benthamiana* (Fig S2B). We therefore speculate that RxL21 co-localisation with DAPI is indicative of an effector mechanism involving transcriptional repression via interaction with TPL.

Consequently, we performed RNA-sequencing to compare gene expression in HA:21 lines with those expressing the truncated form of the effector that is unable to interact with TPL/TPR1 (HA:21ΔEAR lines). This comparison was chosen in order to specifically determine whether RxL21 interaction with TPL/TPR1 is driving alterations in the host transcriptome and eliminate transcriptional effects due to the presence of other domains of the effector. In order to capture transcriptional differences which only arise due to a modified defence response, we induced PTI by treatment with flg22; a conserved stretch of 22 amino acids from bacterial flagellin that is recognised by the receptor FLS2 (Felix, Duran, Volko, & Boller, 1999). We compared two independent HA:21 lines with two HA:21ΔEAR lines, 2 h after both mock treatment and flg22 induction (full details of lines and treatments used for each sample is included in Table S2). The total number of 75 bp paired-end reads generated was 676,059,072 across 24 samples. Transcript abundance was calculated by pseudoalignment of reads to AtRTD2 (Arabidopsis reference transcript dataset) (R. Zhang et al., 2016) using Kallisto (Bray, Pimentel, Melsted, & Pachter, 2016). Genes with low expression across all samples were removed and TMM normalization was performed (Robinson & Oshlack, 2010). RNA count data both before and after filtering and normalisation is provided in Sheets B and C respectively in Table S2.

A PCA plot was used to visualise the data quality and variance between conditions, using the average read counts of biological replicates (Fig S10); samples from independent transgenic lines (HA:21#1/2 or HA:21ΔEAR#1/2) were treated as biological replicates. 49% of variance between the samples can be explained by PC1 (flg22 treatment) however separation between RxL21 and RxL21ΔEAR lines is also observed. Differentially expressed genes (DEGs) were defined as having a log_2_ fold change of 1 or greater, with an adjusted p-value cut-off (after false discovery rate correction) of 0.05. DEGs are listed in Table S3; the total number of DEGs for each comparison is shown in Fig 5A.

**Figure 5.**
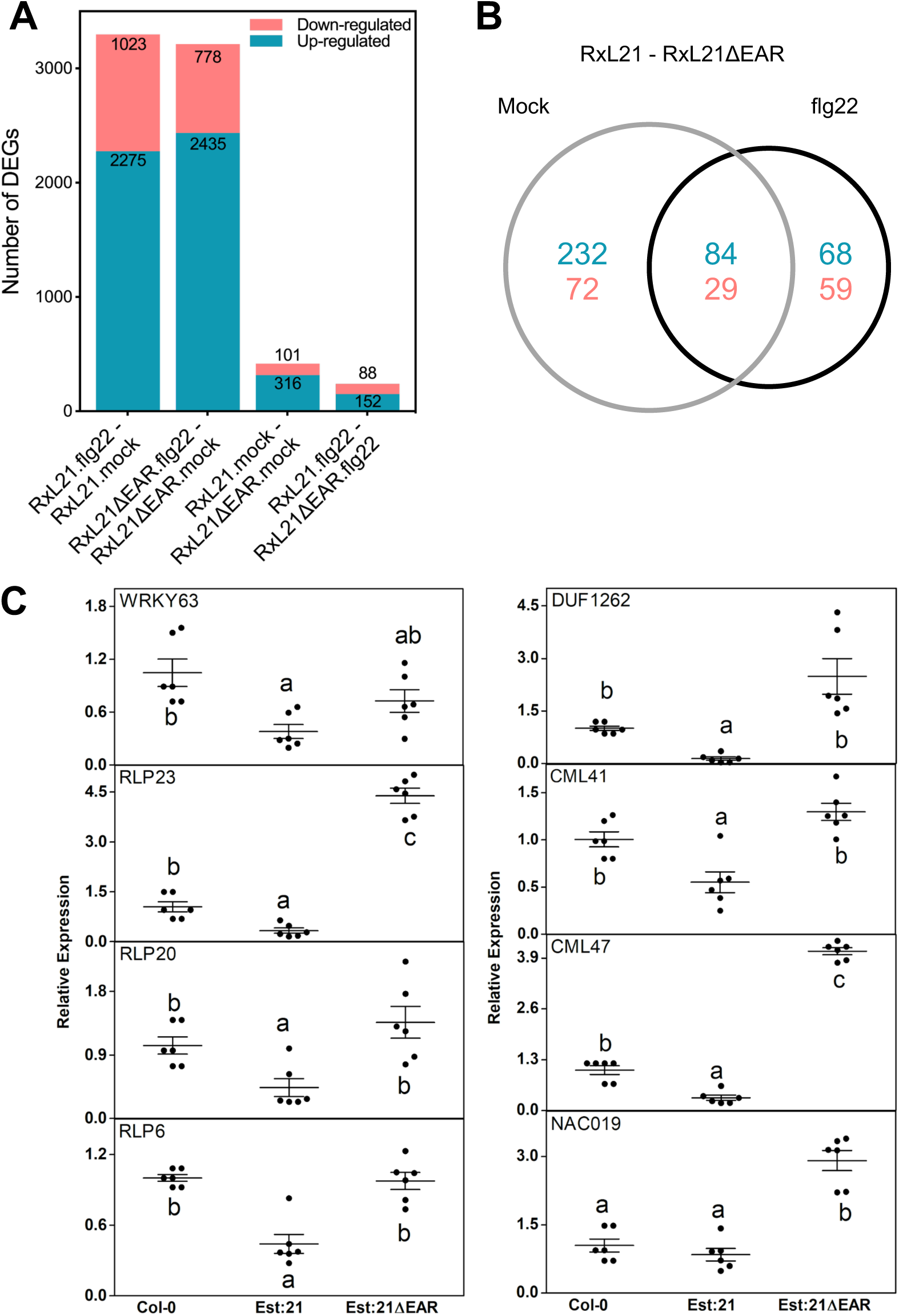
RNAseq identifies differentially expressed genes in RxL21 compared to RxL21ΔEAR transgenic lines. RNAseq was performed on HA-tagged RxL21 and RxL21ΔEAR-expressing transgenic lines. (A) The number of up- and down-regulated genes among differentially expressed genes under mock and flg22 treatment. DEGs were defined as having a log_2_ fold change ≥ 1 or ≤-1, and a BH adjusted p-value of < 0.05. (B) Venn diagram shows differentially expressed genes between RxL21 and RxL21ΔEAR after mock and flg22 treatment with an overlap of 84 upregulated and 29 down regulated genes. (C) Eight genes were selected for validation in estradiol (Est) inducible RxL21 and RxL21ΔEAR expressing lines and Col-0 WT. We treated Arabidopsis seedlings with 30 μM of β-estradiol for 18 hours. For Est:21 and Est:21ΔEAR, data were obtained from 2 independent lines each with 3 biological replicates and expression was normalised to Arabidopsis tubulin gene. Black circles are individual data points and bars denote the mean ± SE of target gene expression. Letters indicate significant differences (P < 0.05) (One-way ANOVA with Tukey’s honest significance difference).

### RxL21 does not perturb the overall PTI response

The scale of transcriptional reprogramming in response to flg22 treatment is similar in both the HA:21 and HA:21ΔEAR lines, and indeed the vast majority of the flg22-induced DEGs are conserved across the two genotypes (2681 genes, with direction of differential expression also conserved: 591 downregulated and 2090 upregulated). Furthermore, DEGs induced upon flg22 treatment in HA:21 and HA:21ΔEAR lines are strongly positively correlated with DEGs in Col-0 at 120 minutes post-flg22 treatment (compared to water treatment) from Rallapalli et al. (2014) (Table S4; Pearson correlation coefficients of 0.841 and 0.844 respectively). This demonstrates that the majority of the flg22 response is intact in HA:21 lines and that RxL21-induced pathogen susceptibility is not due to large-scale interference with the PTI response.

### RxL21 causes differential expression of defence-related genes

There were only 417 genes differentially expressed (DE) between RxL21 and RxL21ΔEAR transgenic lines under mock conditions, and 240 after flg22 treatment (Fig 5A; Sheet B in Table S2). 113 genes were DE in both of these conditions, with the direction of differential expression conserved (Fig 5B). We analysed the DEGs in two groups: those apparent under mock conditions (417 genes, 316 up and 101 down) and the subset of DEGs specific to flg22 treatment (127; 68 up and 59 down) (Fig 5B). To investigate whether transcriptional change in these genes is important for plant immune responses, we first looked for enrichment of Gene Ontology (GO) terms. In both the genes downregulated and upregulated by RxL21 compared to RxL21ΔEAR (under mock conditions) we found significant over-representation of genes involved in response to biotic stimulus/defence and secondary metabolism. Enzymatic activity including oxidoreductase and hydrolase activity, and components of the endomembrane system were also enriched across the DEGs in both mock and flg22 treatments (Table S5).

A significant proportion of the genes whose expression is perturbed by the RxL21 effector are associated with plant immunity (Table S6). 82 genes differentially expressed in response to flg22 in WT plants (Rallapalli et al., 2014) are mis-regulated by the effector, either before or after flg22 treatment. 213 genes differentially expressed in response to RxL21 are differentially expressed during *Pst* infection (DC3000 and/or DC3000HrpA compared to mock) (Lewis et al., 2015), and 142 during *B*. *cinerea* infection (Windram et al., 2012).

We could identify specific cases where RxL21 appeared to be suppressing the host defence response. Fifteen genes that are induced in response to flg22 treatment in WT Arabidopsis (Rallapalli et al., 2014) were downregulated by the full-length effector compared to control after flg22 induction (including *AVRPPHB SUSCEPTIBLE 3, CRK24, CRK38* and *MYB85*). The *Pst* data set profiles gene expression in both virulent *Pst* DC3000 (capable of secreting effector proteins directly into plant cells via the Type III secretion system) as well as a strain DC3000HrpA-which lacks the Type III secretion system. Hence, we can distinguish between host response and pathogen manipulation of gene expression via effector proteins. 20 defence genes that are suppressed by *Pst* effectors were regulated in the same manner by RxL21 (i.e. both *Pst* effectors and RxL21 downregulated or upregulated expression compared to respective controls). Furthermore, 18 genes that are specifically expressed in response to *Pst* effectors (and not part of the host PTI response) were also differentially expressed in response to RxL21. These data indicate not only that RxL21 DEGs are involved in plant immunity, but also that shared mechanisms of host manipulation exist between RxL21 from *Hpa* and *Pst* effectors.

To further strengthen the evidence that RxL21 is altering host gene expression and validate our RNA-seq data, we quantified expression of 8 DEGs using qPCR in an independent set of lines; estradiol-inducible lines Est:21, Est:21ΔEAR and Col-0 control. We selected eight of the genes downregulated by RxL21 compared to RxL21ΔEAR; these genes code for two TFs (WRKY63 and NAC019), three receptor-like proteins, two calmodulins and a protein of unknown function (Sheet B in Table S6). Seven out of the eight genes were significantly downregulated in Est:21 compared to Est:21ΔEAR lines (Fig 5C). Six of these genes also showed reduced expression in Est:21 compared to wildtype Col-0. NAC019 did not differ between Col-0 and Est:21 but was reduced in Est:21 compared to Est:21ΔEAR, and WRKY63 showed an intermediate expression level in Est:21ΔEAR between Est:21 and Col-0. In general the qPCR shows a similar pattern of log_2_ fold change for the 8 selected genes between RxL21 and RxL21ΔEAR lines compared to RNA-seq results (Fig S11).

### RxL21-repressed genes are enriched for TPR1 binding targets

We hypothesised that RxL21 modulates host gene expression via interaction with TPL/TPR1. RxL21 could recruit these corepressors to new sites on the genome (by either binding to the DNA itself or binding to TFs), or RxL21 may maintain TPL/TPR1 repression when it would normally be relieved during infection. Using iDNA-Prot (Lin, Fang, Xiao, & Chou, 2011) we found no evidence to suggest that RxL21 contains a DNA binding motif and hence it is unlikely to bind DNA directly. Next, we looked for enrichment of TF binding motifs in the promoters of the RxL21 DEGs. Promoters of DEGs under mock conditions were significantly enriched for WRKY TF binding motifs in both up and down-regulated genes (Table S7). In the DEGs only evident after flg22 treatment, we found significant enrichment for MYB TF binding motifs (as well as a WRKY binding motif) in the upregulated genes and CAMTA (Calmodulin-binding transcription activator) binding motifs in the genes downregulated by RxL21 (Table S7). This suggests that RxL21 may exert at least some of its effects via modulating TPL/TPR interaction with WRKY, MYB and CAMTA TFs.

We also compared DEGs from our RNAseq analysis to chromatin immunoprecipitation (ChIP)-seq data revealing TPR1 target/bound genes from *pTPR1:TPR1:GFP* Col plants (Griebel et al. (2020), Figure 6). We found that only genes repressed by RxL21 compared to RxL21ΔEAR (both under mock and flg22 treatment) show enrichment for TPR1 target genes (Sheet A, Table S8) with binding observed immediately upstream of the transcription start site (TSS) (Fig 6, purple and orange lines). Hence, for at least some of the RxL21 DEGs, RxL21 appears to impair expression by interaction with TPR1 upstream of the TSS. As expected, the level of TPR1 ChIP signal in RxL21-repressed DEGs is lower than in the group of genes all defined as TPR1 targets from the ChIP-seq data. The 20 genes repressed by RxL21 and identified as TPR1 targets encode several known positive regulators of immunity against biotrophic pathogens (*PBS3, ICS1* and *RLP20*) and regulators of abiotic stress tolerance (*WRKY46, WRKY63, NAC019*) (Table S8). The identification of *WRKY46* and *63* as likely direct targets of RxL21 suppression via TPR1, and the enrichment of WRKY binding motifs in the promoters of RxL21 DEGs suggests that these two TFs could be one key mechanism underlying RxL21-induced pathogen susceptibility. Another key mechanism appears to be mis-regulation of salicylic acid (SA) signalling with RxL21/TPR1 targets including two enzymes required for SA accumulation (ICS1 and PBS3).

**Figure 6.**
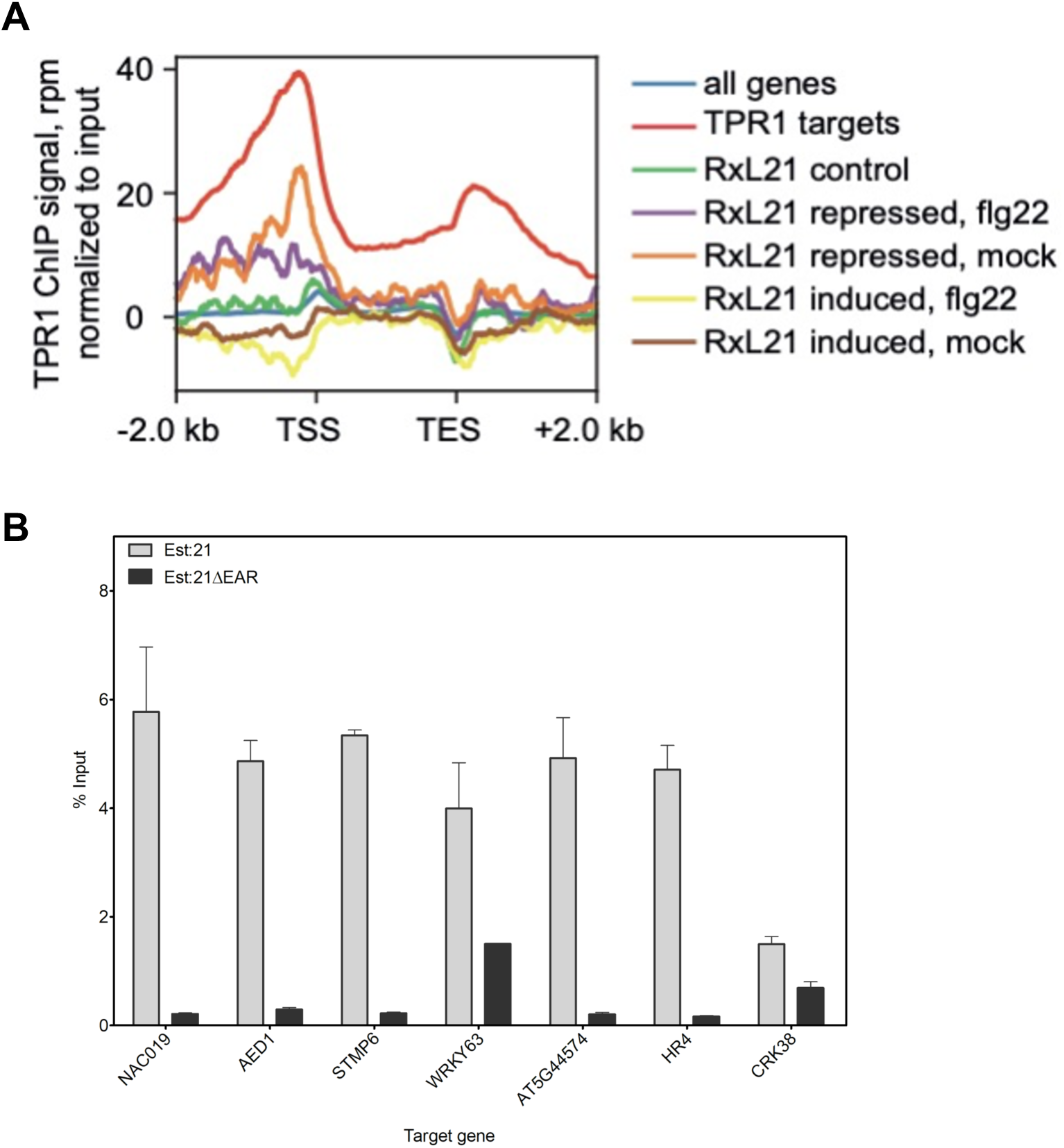
Genes that are bound by TPR1 are overrepresented in genes repressed by RxL21. (A) Arabidopsis genes with repressed expression in HA:RxL21 lines compared to HA:21ΔEAR lines show weak TPR1 binding. Metaplots display the TPR1 enrichment around the transcription start site (TSS) on genes regulated by RxL21 (BH adjusted p-value < 0.05 and log2-fold-change ≥1, with or without flg22) or control genes without evidence for RxL21 dependent expression regulation (RxL21 control). TPR1 bound genes defined in Griebel et al. (2020) were used as a positive control (red line). On the y-axis is mean read count for the TRP1-GFP ChIP samples normalized to the input samples. TPR1 ChIP and input samples were scaled based on the number of mapped reads. TES = transcription end site. (B) ChIP-qPCR of RxL21-repressed genes. Two-week old seedlings, overexpressing RxL21 or RxL21ΔEAR with an N-terminal myc tag and under the control of an estradiol inducible promoter, were treated with 30 μM β-estradiol to induce *RxL21* expression 18 hrs before harvesting and cross-linking with 1% formaldehyde. ChIP assays were performed with an anti-Myc antibody. In the ChIP–qPCR, the enrichment of immunoprecipitated DNA was normalized by the percent input method (signals obtained from ChIP samples were divided by signals obtained from an input sample). Error bar represents ± SE of four technical repeats. Arabidopsis *Actin 2* was included as a control but no amplification was detected after 40 cycles. The experiment was repeated with similar results.

Genes that do not respond to the effector (RxL21 control in Fig 6; Sheet B in Table S8) do not show enrichment for TPR1 binding, while interestingly, genes induced by RxL21 (compared to RxL21ΔEAR in mock or after flg22 treatment; Fig 6, yellow and brown lines) demonstrate depletion of TPR1 binding around the TSS. This suggests that promoters of genes upregulated by RxL21 are not TPR1 targets and hence mis-regulation of these by the effector is likely to be an indirect effect rather than through mis-placing bound TPR1.

To confirm the observed overlap of RxL21 repressed genes with TPR1 binding sites, we performed independent ChIP-qPCR on Est:21 and Est:21ΔEAR lines (Fig 6B) on seven genes (marked in bold in Table S8). Promoter regions encompassing 500 bp upstream and 500 bp down-stream of the transcriptional start site for 5 of these genes (*NAC019, AED1, STMP6, AT5G44574* and *HR4*) were enriched approximately 5-fold in RxL21 immunoprecipitated samples compared to RxL21ΔEAR samples. The promoter regions of *CRK38* and *WRKY63* didn’t show strong enrichment, however we noted that *CRK38* was only repressed by RxL21 after flg22 treatment in our data set. We also amplified *AtActin2* as a negative control which was not detectable in ChIP samples after 40 qPCR cycles. Promoter regions of these 5 genes can, therefore, be directly bound by both RxL21 (and not RxL21ΔEAR) and TPR1. Furthermore, the lack of enrichment in the ChIP-qPCR assays in RxL21ΔEAR samples suggests that RxL21 binds to these promoter regions via TPR1 (maintaining repression when it would normally be relieved) rather than recruiting TPR1 to new locations.

## Discussion

### RxL21 mimics a host gene regulation mechanism

Manipulation of host transcription by effector proteins is a key mechanism by which plant pathogens are able to render the plant a more hospitable environment for colonization. In this study we provide evidence that an effector from the oomycete pathogen *Hpa* mimics a mechanism by which plants regulate transcription throughout their development and in response to stress; the EAR motif is a repression motif that occurs in plant transcriptional repressors and mediates interaction with members of the TPL/TPR corepressor family. This mechanism of TPL recruitment to transcriptional repression complexes is highly conserved across the plant kingdom (Causier, Lloyd, Stevens, & Davies, 2012b; Kagale, Links, & Rozwadowski, 2010; Szemenyei et al., 2008). The *Hpa* effector RxL21 contains a C-terminal EAR motif which we have shown to be responsible for interaction with the CTLH domain of the transcriptional corepressor TPL in both *in vitro* yeast assays and *in planta* (Co-IP and Split YFP; Figure 2 and S3). This interaction is specific, and is abolished even when Leu residues in the EAR motif are substituted with the similar amino acid Ile. A pivotal role for TPL family members in regulation of immune responses has been previously implicated (Robert-Seilaniantz, Grant, & Jones, 2011; Zhu et al., 2010) and Arabidopsis plants lacking expression of TPL and its closest homologues, TPR1 and TPR4, show enhanced susceptibility to *Pst, Hpa* and *B*. *cinerea* (Zhu et al., 2010 and Figure 3). Interestingly, the effector RxL21 also interacts with TPR1 and, when expressed *in planta*, enhances susceptibility of Arabidopsis to a biotrophic (*Hpa*), hemibiotrophic (*Pst*) and necrotrophic pathogen (*B*. *cinerea*). There are a few examples of EAR motifs in other pathogen effectors - the bacterial XopD effector family (J.-G. Kim et al., 2011) and an effector from fungi that contains an EAR motif and interacts with TPR4 (Petre et al., 2015). Crucially we show here that the C-terminal EAR motif (and thus interaction with TPL/TPR1) is required for enhanced pathogen susceptibility provided by the RxL21 effector (Figure 4).

### RxL21 alters expression of a subset of the host defence response

By performing RNA-seq comparing the transcriptional effects of expressing RxL21 compared to the effector lacking the EAR motif (RxL21ΔEAR) we were able to identify specific DEGs that result from RxL21-TPL/TPR1 interaction. Importantly, there is no large-scale change in gene expression in response to presence of the effector (either with or without flg22 treatment to induce PTI), yet effector expression still delivers a striking susceptibility enhancement to the host plant. This is similar to the situation with *Pst* where blocking of effector delivery through the Type III Secretion System only perturbed the expression of 872 PTI-regulated genes with the vast majority (5350) showing no change in expression or amplification of usual PTI-regulation (Lewis et al., 2015). These findings highlight the ability of pathogen effectors to target (and hence identify) specific components of the plant defence response with great effect.

The genes mis-regulated by RxL21 include several important components of plant defence responses. These include down regulation of several receptor-like proteins (RLPs); *RLP6, RLP22, RLP23* and *RLP35*, with *RLP20* DE specifically after flg22 treatment. RLPs are the second largest family of Arabidopsis leucine rich repeat-containing receptors; several members have been shown to function as PAMP receptors and it is becoming increasingly clear that RLPs play a critical role in pathogen recognition (Jamieson, Shan, & He, 2018). Consistent with this, many RLPs (including *RLP6, 22, 23* and *35*) show increased expression during pathogen infection and/or hormone treatment (Jamieson et al., 2018; J. Wu et al., 2016) and several RLPs are involved in fungal and oomycete resistance (Albert et al., 2015; Jiang et al., 2013; Ramonell et al., 2005; Shen & Diener, 2013; W. Zhang et al., 2013). In addition, several other genes downregulated by RxL21 are known components of the defence response. These include genes that impact resistance against biotrophic pathogens such as *accelerated cell death 6* (*ACD6*) and Cysteine-rich receptor-like kinase *CRK13* (Acharya et al., 2007) that positively regulate SA signalling (Todesco et al., 2010), and Calmodulin-like *CML41* that regulates flg22-induced stomatal closure (Xu et al., 2017). Furthermore, miR825 target *AT3G04220* (Nie et al., 2019) and the lipid transfer protein AZI3 (Chassot, Nawrath, & Métraux, 2007) contribute to defence against necrotrophs and the lectin receptor kinase LecRK4.1 positively regulates Arabidopsis PTI and contributes to resistance against both biotrophic and necrotrophic pathogens (Singh et al., 2012). Reduction in expression of these genes is likely to contribute to the observed enhanced susceptibility of RxL21 over-expressing plants to both biotrophic and necrotrophic pathogens.

Twenty genes that are direct TPR1 targets are repressed by RxL21, seven of which were specific to flg22 treatment. This includes tetraspanin9 (*TET9*) which was 2.6-fold repressed by RxL21 compared to RxL21ΔEAR. TET8/9 are involved in the formation of exosome-like extracellular vesicles to deliver host sRNAs into *B*. *cinerea* to decrease fungal virulence (Cai et al., 2018). TET9 accumulates around fungal infection sites after *B*. *cinerea* infection and *tet9* loss-of-function mutants display weak but consistent enhanced susceptibility towards *B*. *cinerea* (Cai et al., 2018), leading us to speculate that *TET9* repression could contribute to the *B*. *cinerea* susceptibility we see in RxL21-expressing lines.

Two WRKY TFs (WRKY46 and WRKY63) are direct TPR1 targets and repressed in RxL21 expressing plants in comparison to the RxL21ΔEAR variant. WRKY TFs have been shown to regulate defence responses against biotrophic and necrotrophic pathogens (Birkenbihl, Diezel, & Somssich, 2012; S. Liu, Ziegler, Zeier, Birkenbihl, & Somssich, 2017) and WRKY46 overexpressing plants are more resistant to *Pst* (Hu, Dong, & Yu, 2012). Moreover, non-host resistance against *Erwinia amylovora* is regulated by WRKY46 and WRKY54 via EDS1 (Moreau et al., 2012) and WRKY46 is a transcriptional activator of PBS3 expression (van Verk, Bol, & Linthorst, 2011). Consistent with this, *PBS3* expression is also reduced in RxL21 compared to RxL21ΔEAR over-expressing plants. Recently, it was shown that PBS3 protects EDS1 from proteasomal digestion (Chang et al., 2019), hence reductions in PBS3 expression could lead to lower levels of EDS1 protein and enhanced susceptibility to biotrophic pathogens (Parker et al., 1996). Both WRKY46 and 63 bind to the promoter of the SA biosynthesis gene *SID2* with increased levels of SA correlated with increased levels of expression of these and several other WRKY TFs (S. Zhang et al., 2017). Consistent with this, *SID2* (also called *ICS1*) is downregulated by RxL21 after flg22 treatment, however this may also be due to direct repression via association with TPR1 as *SID2* is also a target of TPR1.

WRKY TF binding motifs were overrepresented in the promoters of RxL21 DEGs implying that many of the DEGs are downstream targets of WRKY TFs. It is possible that WRKY46 and WRKY63 are directly repressed by RxL21 which in turn leads to changes in expression of their target genes. We observed overrepresentation of CAMTA motifs in genes that were specifically downregulated by RxL21 in an EAR-dependent manner upon flg22 treatment, indicating that target genes of CAMTA TFs are showing differential expression. There is no evidence to date that CAMTA TFs are TPL/TPR targets (Causier, Ashworth, Guo, & Davies, 2012a) but CAMTA TFs play a role in immune regulation through suppressing pathogen-responsive genes and could therefore be an important target for RxL21 manipulation via TPL/TPR1 (Jacob et al., 2018; Y. Kim, Gilmour, Chao, Park, & Thomashow, 2020; Yuan, Du, & Poovaiah, 2018).

### Recruitment of RxL21 to TPL/TPR1 transcriptional complexes

We have shown that the suppression of immunity by RxL21 depends on its EAR domain, and hence most likely through modifying actions of TPL and TPR1. Several scenarios are possible. On the one hand, RxL21 could interact with a TF and mediate subsequent TPL/TPR1 binding (to novel sites) in a manner similar to NINJA or JAZ proteins (Pauwels et al., 2010). Evidence for an interaction of RxL21 with the TF TCP14 was seen in Y2H experiments (Mukhtar et al., 2011) and later it was shown that TCP14 shifts RxL21 into subnuclear foci (Weßling et al., 2014). However, if RxL21 was binding TFs independently of TPL/TPR1 then our ChIP-PCR experiments would be expected to show a similar enrichment in the RxL21 and RxL21ΔEAR at the tested loci. An alternative model is that RxL21 interferes with the repression (or lifting of repression) of existing TPL/TPR1 targets. We provide evidence to show that, at least for some genes, RxL21 appears to bind to TPL/TPR1 within transcriptional complexes at plant gene promoters and prevent transcriptional de-repression. Firstly, there is a significant over-representation in TPR1 binding sites upstream of genes that are repressed by full-length RxL21 compared to RxL21ΔEAR (where interaction with TPL/TPR1 is lost by deletion of the EAR motif). Secondly, promoters of several RxL21-repressed genes were shown to be not just binding targets of TPR1 but also binding targets of RxL21 (and not of RxL21ΔEAR). This indicates that in at least these cases, RxL21 likely binds to TPR1 protein already bound at the gene promoter (as there is no binding of RxL21 to these promoters without the EAR motif) and subsequently exerts its activity on the TPL/TPR1 complex.

The TPL family of proteins are closely related to Groucho / Tup1 proteins in animals and fungi (Z. Liu & Karmarkar, 2008) which, like TPL, have been shown to form tetrameric structures. Tetramerization has been suggested as a mechanism to enable recruitment of multiple TFs to a single complex (Martin-Arevalillo et al., 2017) and binding of EAR motif peptides does not prevent tetramerization (G. Chen, Nguyen, & Courey, 1998; Martin-Arevalillo et al., 2017; Nuthall, Husain, McLarren, & Stifani, 2002). It is therefore plausible that RxL21 is able to bind TPL/TPR1 via its EAR motif while other epitopes of the TPL oligomer are binding other proteins such as TFs within the transcriptional complex. How TPL/TPR1-bound RxL21 behaves is not known. However, TPR1 activity was shown to be regulated by the (SUMO) E3 ligase SIZ1 (Niu et al., 2019), perhaps via SUMOylation inhibiting the corepressor activity of TPR1 by preventing its interaction with HISTONE DEACETYLASE 19 (HDA19). It is possible that RxL21 is shielding the SUMO attachment sites K282 and K721 in TPR1 and/or preventing (SUMO) E3 ligase activity, thereby enhancing the TPL/TPR co-repressor activity.

There is no evidence that the only mechanism of RxL21 action is through maintained repression of direct TPL/TPR1 targets and it is worth noting that many DEGs were upregulated in the presence of RxL21. In addition to immune suppression, pathogen effectors are known to directly manipulate plant gene expression, and hence physiology, to aid infection (L.-Q. Chen et al., 2010; Fatima & Senthil-Kumar, 2015). During *Pst* infection, more than 1500 genes were specifically upregulated in response to *Pst* effectors, and we did observe an overlap (15 genes) between these genes and those upregulated by RxL21. However, it is possible that upregulated genes could be downstream targets of TFs or other regulators targeted by RxL21. This appears to be the most likely explanation given that the RxL21 upregulated genes are not enriched for TPR1 binding in wildtype plants and the enrichment for WRKY TF binding motifs in the promoter regions of these genes.

Remarkably, the pathogen effector RxL21 alone can increase susceptibility to pathogens with varying lifestyle and virulence strategies. To our knowledge there are very few, if any, effectors that exhibit this activity. RxL21 is one of several effectors that mimic the EAR motif for transcriptional repression, and appears to actively initiate and/or maintain repression of gene expression mediated by TPL/TPR1. We show here that the RxL21 EAR motif is essential for its virulence function, and for modifying the expression of key host defence genes. Future interrogations will be to examine the RxL21 mode of action on TPL/TPR1 transcriptional repression complexes and determine which effector-manipulated defences ultimately result in enhanced susceptibility of the host plant.

## Methods

### Sequence alignment

Alignment of HaRxL21 alleles was performed using sequences in Asai et al. (2018) or BioProject PRJNA298674 (for Noks1). Alignment was performed using T-COFFEE (Version_11.00.d625267; http://tcoffee.crg.cat/apps/tcoffee/do:mcoffee). SignalP-5.0 (http://www.cbs.dtu.dk/services/SignalP/) was used to predict the signal peptide cleavage site.

### Yeast-2-Hybrid

Yeast-2-Hybrid (Y2H) screening was performed as described in Dreze et al. (2010). Briefly, yeast strains Y8800 and Y8930 harbouring AD-X and DB-X constructs respectively were mated on yeast-extract, peptone, dextrose, Adenine (YPDA) medium. Yeast was replica plated onto Synthetic complete (SC) media lacking combinations of Leu, Trp, His and Ade. 3AT (3-amino-1,2,4-triazole) was added to selective plates lacking His to increase stringency as a competitive inhibitor of the HIS3 gene product. Plates were cleaned using sterile velvets after 1 day and imaged after 4 days. To generate EAR motif mutants, site-directed mutagenesis (SDM) was performed using a QuikChange II Site-Directed Mutagenesis Kit (Agilent, Santa Clara, US). Primer sequences for SDM are included in Table S9.

### Plant material

35S::HaRxL21 lines (RxL21a/b) are described in Fabro et al. (2011). To generate 35S::HA::HaRxL21 lines in Arabidopsis, RxL21 and RxL21ΔEAR were cloned into pEarleygate201 (Earley, Haag, Pontes, & Opper, 2006) and transformed into Arabidopsis ecotype Col-4 using floral dipping {Clough:1998vw}. Independent transformants were selected on BASTA (Bayer CropScience, Wolfenbüttel, Germany) until homozygous. Western Blotting was performed on 14 day old seedlings to determine protein expression using anti-HA high affinity antibody (Roche, Penzberg, Germany). Primer sequences for RxL21 and RxL21 deletion variants are in Table S9. To generate estradiol inducible lines, RxL21 and RxL21ΔEAR were cloned into the pER8 plasmid {Zuo:2000} and transformed into Arabidopsis (Col-0 background) via floral dip. Independent homozygous lines were selected on hygromycin (Invitrogen, Carlsbad, US).

### Pathogen Assays

*Hpa* infection screens were performed as described in Tomé et al. (2014). Briefly, plants were sown in a randomized block design within the inner modules of p40 seed trays at a density of approximately 30 seedlings per module. Modules at the periphery of the tray were sown with WT. Plants were grown under a 12 h photoperiod, 20°C, 60% humidity. *Hpa* isolates Noks1 and Maks9 were maintained on Col-0 plants and inoculated onto 2 week old seedlings at 3×10^4^ spores/mL. After infection, plants were covered with a lid and sealed. Sporangiophore counts were performed using a dissecting microscope at 4 dpi. Botrytis infection screens were performed as described in Windram et al. (2012). Briefly, *B*. *cinerea* (strain pepper) was maintained on sterile tinned apricot halves in petri dishes, kept in the dark at 25°C. Detached Arabidopsis leaves were placed on 0.8% agar and inoculated with a 10 μL droplet of spore suspension at 1×10^5^ spores/mL in 50% sterile grape juice. Lesions were imaged at 24, 48, 72 and 96 h post infection (hpi). For Botrytis infections on estradiol-inducible lines, plants were grown at 10 h photoperiod, 16°C at 60% humidity. Gene expression was induced by 30 μM β-estradiol, 18 h before infection. Spores were washed from the surface of the *B*. *cinerea* culture plate using potato dextrose broth (PDB) medium and leaves were infected using a 5 μL droplet at a concentration of 1×10^4^ spores/mL.

### Localisation and Split YFP

For localisation, pK7WGF2 (N-terminal GFP fusion; (Karimi, Inzé, & Depicker, 2002)) and pB7WGR2 (N-terminal RFP fusion; (Karimi, De Meyer, & Hilson, 2005)) vectors were used. BiFP (Bimolecular Fluorescence complementation in Planta) vectors were used for generation of C- or N-YFP fusion constructs (Azimzadeh et al., 2008). Expression vectors were transformed into *A*. *tumefaciens* strain GV3101 and cultured overnight at 28°C. *A*. *tumefaciens* harbouring expression constructs, along with P19 (Voinnet, Rivas, Mestre, & Baulcombe, 2003) were co-infiltrated into *N*. *benthamiana*. Leaves were imaged 48 h after infiltration by laser scanning confocal microscopy.

### Co-immunoprecipitation

For Co-immunoprecipitation (Co-IP), TPL and RxL21 were expressed in pCsVMV-HA3-N-1300 (C-terminal HA tag) and Per8 (N-terminal myc tag) vectors. Proteins were expressed transiently in *N*. *benthamiana*. Leaves of about four-week-old plants were infiltrated with *A*. *tumefaciens* (OD600 = 0.2) in infiltration buffer (10 mM MgCl2, 10 mM MES, and 100 μM acetosyringone). After 24 hpi, leaves were sprayed with 30 μM estradiol to induce the expression of RxL21 and subsequent mutants. At 48 hpi, approximately 2 g of tissue from the infiltrated area was collected and frozen with liquid N_2_, then ground into powder using a mortar and pestle. All of the steps were carried out either on ice or in a 4°C cold room. About three volumes of nuclei extraction buffer (NEB) (20 mM Hepes pH 8, 250mM Sucrose, 1mM Magnesium Chloride, 5mM Potassium Chloride, 40% (v/v) glycerol, 0.25% Triton X-100, 0.1 mM PMSF, Protease inhibitor cocktail (Roche cOmplete), were added to each sample. The resuspended samples were filtered using miracloth and centrifuged at 3,320 g for 15 min at 4°C. The pellet was subsequently resuspended and washed with NEB in 2 mL microcentrifuge tubes. Washed pellet was resuspended in lysis buffer (10 mM Tris pH7.5, 0.5 mM EDTA, 1 mM PMSF, 1% Protease Inhibitor, 150 mM Sodium Chloride, 0.5% Igepal) and sonicated to break the nuclei. After sonication, the supernatant was centrifuged again to remove additional debris, and the supernatant was used as input for IP. Afterwards, the µMACS kit of magnetic microbeads, conjugated to an anti-c-myc monoclonal antibody (130-091-123; Miltenyi Biotech), was used for IP according to the manufacturer’s instructions. Eluted proteins were analysed by western blot using HRP-conjugated anti-myc antibody (130-092-113; Miltenyi Biotec), or an anti-HA antibody (3F10; Roche).

### RNA-Seq

#### Sample preparation

Arabidopsis seedlings were grown on ½ MS agar with sucrose for 14 days at 22°C with photoperiod of 8 h light and 16 h dark, and 150 mmol photons * m^-2^ × s^-1^. Seedlings were then transferred to 6 well plates containing 5 mL ½ MS liquid media (approximately 8 seedlings per well) and left to rest overnight. Media was replaced with 5 mL fresh ½ MS liquid media containing 100 nM flg22 or 5 mL ½ MS control. Each sample consisted of total pooled seedlings from a well, these were harvested 2 h post induction, briefly dried on tissue and flash frozen in liquid nitrogen. For each of 35S::HA::RxL21 and 35S::HA::RxL21ΔEAR, two independently transformed Arabidopsis lines were used (HA:21#1/2 and HA:21ΔEAR#1/2), each with 3 biological replicates. RNA was extracted from Arabidopsis tissue using Trizol (Invitrogen) and cleaned up using a RNeasy Mini Kit (Qiagen, Hilden, Germany) including on-column DNase treatment using a RNase-free DNase kit (Qiagen). Libraries were made using the NEBNext Ultra Directional RNA library prep kit. Each library was sequenced on two lanes of Illumina HiSeq 4000 generating 75 bp paired end reads. The sequencing files were examined using fastQC version 0.11.5.

#### Quantification of Transcripts and DE Gene Analysis

Transcript abundances were generated using Kallisto version 0.44.0 (Bray et al., 2016) using an index generated using Arabidopsis Thaliana Reference Transcript Dataset 2 (R. Zhang et al., 2016). The number of bootstraps was set to 100. DEGs were generated using the 3D-RNA-seq app (Wenbin Guo et al., 2019). Sequencing replicates from the two HiSeq 4000 lanes were merged. Data was pre-processed by filtering to remove genes which did not meet the following criteria; 1) An expressed transcript must have at least 3 out of 23 samples with CPM (count per million read) ≥1 and 2) An expressed gene must have at least one expressed transcript. One sample was removed from further analysis (RxL21ΔEAR.flg22 line 2, biorep 3) due to outlying sequencing depth. RNA-seq read counts were normalised with Trimmed Mean of M-values method (Robinson & Oshlack, 2010). Models of expression contrasts (RxL21.flg22 vs RxL21.mock, RxL21ΔEAR.flg22 vs RxL21ΔEAR.mock, RxL21.mock vs RxL21ΔEAR.mock and RxL21.flg22 vs RxL21ΔEAR.flg22) were fitted using Limma Voom (Law, Chen, Shi, & Smyth, 2014; Ritchie et al., 2015). Genes were significantly DE in the contrast groups if they had BH adjusted p-value < 0.05 and log_2_-fold-change ≥1. We performed Go-Term analysis for biological process, cellular component and molecular function using AgriGO ((Tian et al., 2017), http://bioinfo.cau.edu.cn/agriGO/). Significance of GO enrichment was determined with FDR < 0.05. Known Arabidopsis TF DNA-binding motifs were retrieved from CIS-DB version 1.02 (Weirauch, Yang, Albu, & Cote, 2014), and those described in Franco-Zorilla et al., (2014). Promoter sequences defined as the 500 bp upstream of the transcription start site were retrieved from https://www.arabidopsis.org/ (Araport 11 annotation). Motif occurrences were determined using FIMO (C. E. Grant, Bailey, & Noble, 2011), and promoters defined as positive for a motif if it had at least one match with a *P* value < 10^−4^. Motif enrichment was assessed using the hypergeometric distribution against the background of all genes. *P* values < 0.001 were considered significant to allow for multiple testing. RNA seq data is available online at NCBI under Bioproject ID: PRJNA622757.

### qPCR

Total RNA extraction was performed with TRIzol (Invitrogen) following the manufacturer’s instructions. cDNA was synthesized by using the RevertAid cDNA synthesis kit (Thermofisher Scientific, Waltham, US). Quantitative PCR was performed in the Applied Biosystems 7500 FAST real-time PCR system using SYBR Green JumpStart Taq ReadyMix (Sigma-Aldrich, St Louis, US). Transcript levels of target genes were determined via the 2-ΔΔCt method (Livak & Schmittgen, 2001), normalized to the amount of *Arabidopsis tubulin 4* (*AT5G44340*) transcript. For expression analysis of HA:RxL21 and HA: 21ΔEAR lines, expression was normalised to *Actin2* (*AT3G18780*) and *UBQ5* (AT3G62250) transcript levels. Primer sequences are detailed in Table S9.

### Chromatin immunoprecipitation (ChIP) PCR

ChIP PCR was performed as described in Gendrel et al. (2005). Briefly, two week old seedlings were harvested and cross-linked in 1% formaldehyde (Sigma-Aldrich) solution under vacuum for 15 min. The isolated chromatin complex was resuspended in Nuclei lysis buffer 50 mM Tris-HCl, 10m M EDTA, % SDS and one tablet PI (Roche cOmplete) and sheared by sonication (Bioruptor, Diagenode, Ougrée, Belgium) to reduce the average DNA fragment size to around 500 bp. The sonicated chromatin complex was diluted in ChIP dilution buffer (16.7 mM Tris-HCl, 1.2 mM EDTA, 167 mM NaCl, 1.1% Triton X-100 and one tablet PI) and immunoprecipitated by anti-myc antibody (ab9132; ChIP-grade, Abcam, Cambridge, UK). After reverse cross-linking, the immunoprecipitated DNA was extracted by using equal amounts of phenol/chloroform/isoamyl alcohol and precipitated with 100% EtOH, 1/10 volume of 3 M Sodium acetate and, 10 mg/mL glycogen. DNA was resuspended in Milli-Q water and analyzed by qPCR with gene specific primers. Primer sequences are shown in Table S9. Input % in IP samples was calculated by (100*2^(Adjusted input-IP)). *AtActin2* (*AT3G18780*) was included as a control.

### Metaplots for TPR1 binding

Preprocessing of chromatin immunoprecipitation-sequencing (ChIP-seq) data for the *pTPR1:TPR1-GFP* Col-0 line (Zhu et al., 2010) was performed as in Griebel et al. (2020). Briefly, adapters and other overrepresented sequences detected with fastqc (version 0.11.9; http://www.bioinformatics.babraham.ac.uk/projects/fastqc/) were removed with cutadapt (version 1.9.1, -e 0.2 -n 2 -m 30; (Martin, 2011)). Raw reads were mapped to *Arabidopsis thaliana* genome version TAIR10 (arabidopsis.org) with bowtie2 (version 2.2.8; (Langmead & Salzberg, 2012)). The alignment files were filtered for low-quality reads (samtools view -q 10, version 1.9, (Li, 2011; Li et al., 2009)), deduplicated and merged from the three biological replicates. The metaplots were prepared with deepTools 3.3.0 following the manual pages (https://github.com/deeptools/deepTools/tree/develop). The TPR1-GFP ChIP-seq data were input-normalized using the bamCompare function (--operation subtract, default ‘readCount’ scaling).

## Acknowledgments

Thank you to Prof. Karl-Heinz Kogel for his support and the opportunity to continue this project; Sally James (York University Biology Technology facility) for performing RNAseq library prep; Tina Payne and Christina Neumann for their assistance with transgenic plants; Dr Hazel McLellan (James Hutton Institute, Dundee) for providing the NbTPL construct; Prof. Jeffrey Long for sharing with us the TPLΔCTLH mutant construct; Dr Barry Causier for sharing TPR1-4 constructs; Dr. Yuelin Zhang for sharing the *tpr1*-*tpl*-*tp4* mutant line; Prof. David Mackey (The Ohio State University, Columbus) for sharing pCsVMV-HA3-N-1300 vector and Dr Francois Parcy (University Grenoble, France) for the BiFP vectors. This work was supported by BBSRC grant BB/K018612/1. SH was supported by the University of York and PK by DAAD scholarship. DL, TG and JEP were supported by the Max-Planck Society and Deutsche Forschungsgemeinschaft (DFG; German Research Foundation) under Germany’s Excellence Strategy CEPLAS (EXC-2048/1, Project 390686111; JEP, DL), CRC 680 project B10 (JEP, DL) and CRC 670 project TP19 (JEP, TG).

## Author Contributions

Conceived and designed the experiments: JS, JB, SH, KD, PK, TG, DL, JEP. Performed the experiments: SH, PK, JS, TG, DL. Analysed the data: SH, PK, JS, KD, RH, WG, RZ, DL, TG, JB, JEP. Wrote and edited the manuscript: SH, PK, JS, and KD with input from DL, TG and JEP.

## Supporting Data

**Table S1. *Hpa* RxLs and RxLLs found to contain the LxLxL motif**. Putative RxL and RxL-like (RxLL) effectors from *Hpa* were searched for the presence of a LxLxL motif. Effectors were characterised as having a C- or N-terminal EAR motif if the motif was detected within 35 amino acids of the N-or C-terminus of the effector sequence (otherwise ‘mid’). Nuclear or cytoplasmic localisation of effectors as identified by Caillaud et al., (2011) is indicated. Effectors shown in Fig S4 and tested for interaction with TPL are highlighted in yellow.

**Table S2. RNA Seq sample information and read counts**. (a) Line and treatment information for each sample. (b) Raw read counts obtained from Kallisto before any processing. Sequencing replicates are indicated by ‘srep’. (c) Normalised read counts per gene after combining sequencing replicates, removing lowly expressed genes across all samples and normalisation using the limma-voom pipeline.

**Table S3. Differentially expressed genes after RNA-seq analysis**. Differentially expressed genes after RNA-seq analysis. Contrast indicates in which pair-wise comparison the gene is differentially expressed. Adj.pval indicates significance after limma-voom pipeline and Benjamini Hochberg false discovery rate correction. FC = log2 fold change. Arabidopsis gene descriptions are based on Araport 11 annotation.

**Table S4. Comparison of flg22 induced gene expression in RxL21 lines compared to Col-0 WT**. Log2 fold expression of differentially expressed genes in RxL21 and RxL21ΔEAR lines after flg22 treatment compared to mock, compared to Log_2_ fold expression in Col-0 at 120 minutes post treatment with flg22 compared to mock treatment from Rallapalli et al. (2014).

**Table S5. Over-represented GO categories in differentially expressed genes between RxL21 and RxL21ΔEAR in both mock and flg22 conditions**. Singular enrichment analysis of GO terms was performed using AgriGO. The cutoff P-value after false discovery rate correction is <0.05. (F = molecular function, P = biological process, C = cellular component).

**Table S6. (A) Genes differentially regulated in RxL21 compared to RxL21ΔEAR**. Gene descriptions are from Araport 11. Adj.pval indicates significance after limma-voom pipeline and Benjamini Hochberg false discovery rate correction. Log_2_ fold change (FC) indicates expression in HA:RxL21 lines compared to HA:RxL21ΔEAR lines. Flg22 specific DEGs only show significant differential expression after flg22 induction. Mock / flg22 independent DEGs are DE under mock and/or mock and flg22 conditions. Flg22 (Column H) indicates expression values 120 minutes after flg22 induction from Rallapalli et al. (2014). Comparison is shown to *B*. *cinerea* responsive DEGs from Windram et al. (2012) and DEGs during *Pst* infection (Lewis et al. 2015), including (Column K) description of *Pst* expression type from Figure 4 (Lewis et al, 2015) where applicable. (B) Details of the genes used for qPCR for RNAseq verification.

**Table S7. Motifs Over-represented in RxL21 differentially expressed genes identified using RNA-seq**. Known Arabidopsis TF DNA-binding motifs were retrieved from CIS-DB version 1.02 (Weirauch et al., 2014), and those described in Franco-Zorrilla et al. (2014). Motif occurrences were determined using FIMO (Grant et al., 2011) and enrichment was assessed using the hypergeometric distribution against the background of all genes. Number of genes indicates genes containing each motif within 500 bp promoter region compared to the total number of genes in each comparison (DEG list). P-value cut off is 0.001.

**Table S8. A) Overlap between genes associated with TPR1 binding sites and genes differentially regulated in Arabidopsis plants expressing the RxL21 effector**. “RxL21 induced” or “RxL21 repressed” indicates genes which show significantly higher or lower expression respectively in HA:21-expressing plants compared to HA:21ΔEAR-expressing plants. Number of genes for each comparison is shown with overlap to TPR1 targets in brackets. P values indicate significance of overlap between each group of genes with TPR1 targets. Genes selected for verification by ChIP-PCR are indicated in bold (Column D). Gene descriptions taken from Araport 11. B) The control set of 150 genes which show no differential expression between HA:21 and HA:21ΔEAR.

**Table S9. Primer sequences used in this study**.

---

**Figure S1.**
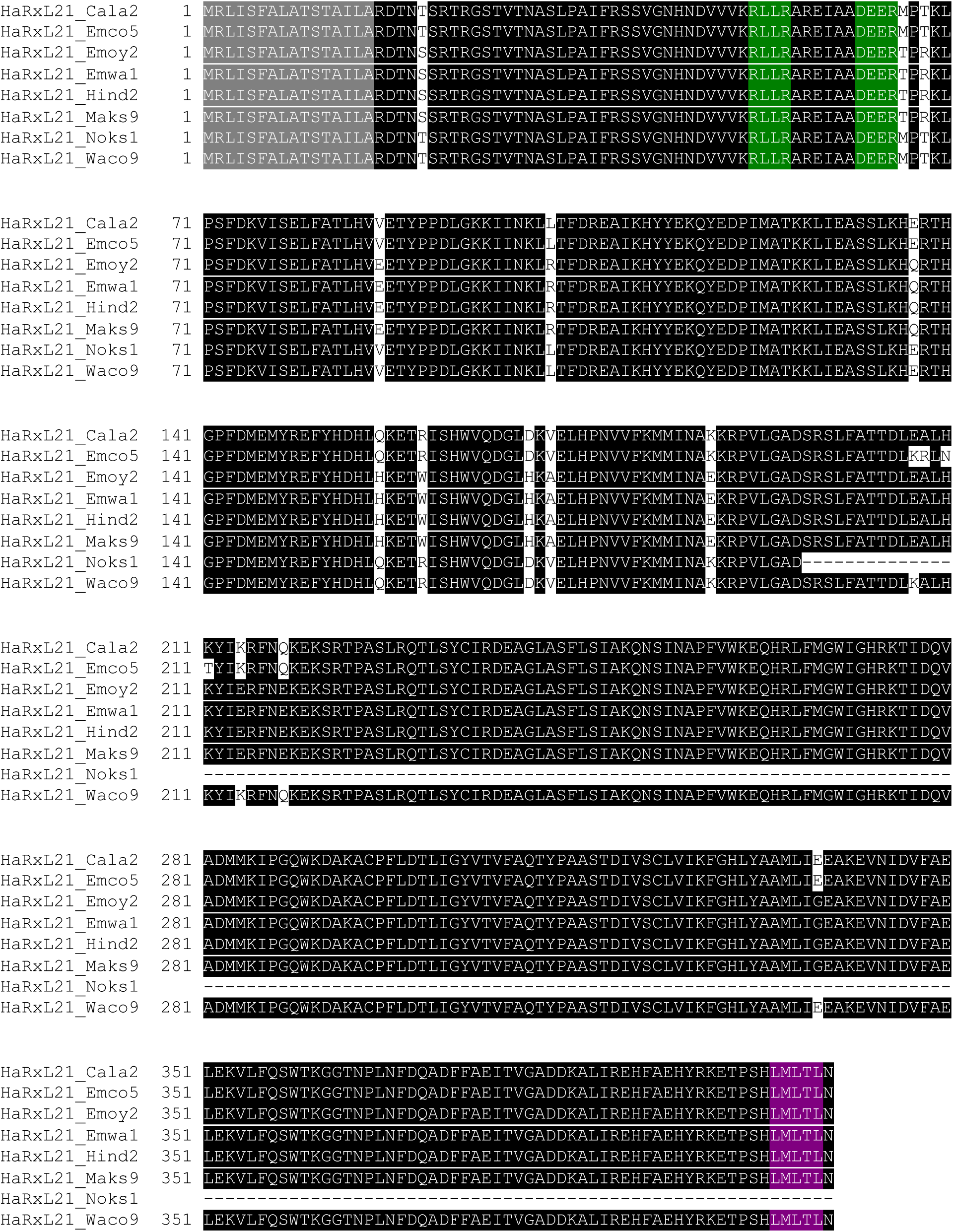
Amino acid sequence alignment of HaRxL21 alleles from *Hpa* isolates Cala2, Emco5, Emoy2, Emwa1, Hind2, Maks9, Noks1 and Waco9. Sequences were obtained from Asai et al., 2018 (Cala2, Emco5, Emoy2, Emwa1, Hind2, Maks9 and Waco9) and BioProject PRJNA298674 (Noks1). Multiple sequence alignment was performed using T-coffee (http://tcoffee.crg.cat/apps/tcoffee/do:mcoffee). Sites of amino acid substitution between alleles are highlighted in white. Predicted signal peptide is shown in grey. The RxLR-DEER motif (green) and EAR motif (magenta) are conserved across alleles except Noks1.

**Figure S2.**
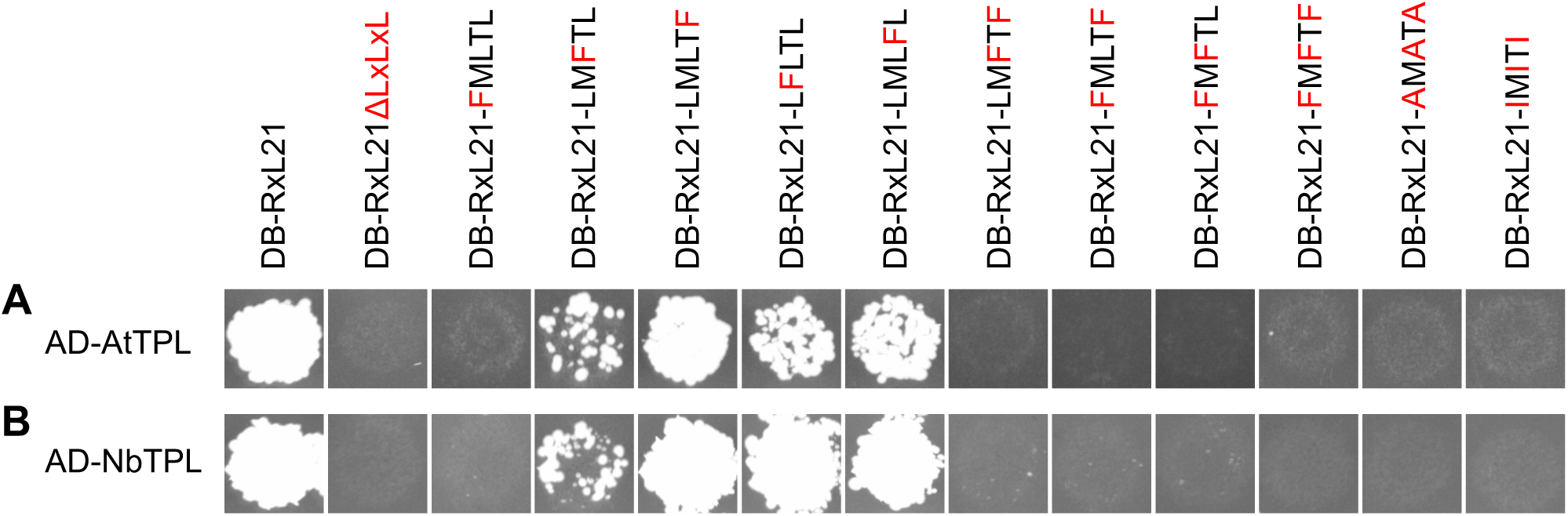
Site directed mutagenesis of Leucine residues in the EAR motif of RxL21 abolishes interaction with TPL by Yeast-2-Hybrid. Leu residues in the EAR (LxLxL) motif were mutated to Phenylalanine (F), Alanine (A) and Isoleucine (I). Amino acids that differ from WT are indicated in Red. AD; GAL4 activation domain. DB; GAL4 DNA binding domain. (A) Interaction was tested against TPL from Arabidopsis (AtTPL). (B) Interaction was tested with TPL from *N*. *benthamiana* (NbTPL). Growth on media lacking Leu, Trp and His is shown, indicating successful mating and activation of the *GAL1::HIS3* reporter gene due to interaction. All combinations tested showed growth on media lacking only Leu and Trp indicating successful mating (not shown). The experiment was repeated on multiple plates with similar results.

**Figure S3.**
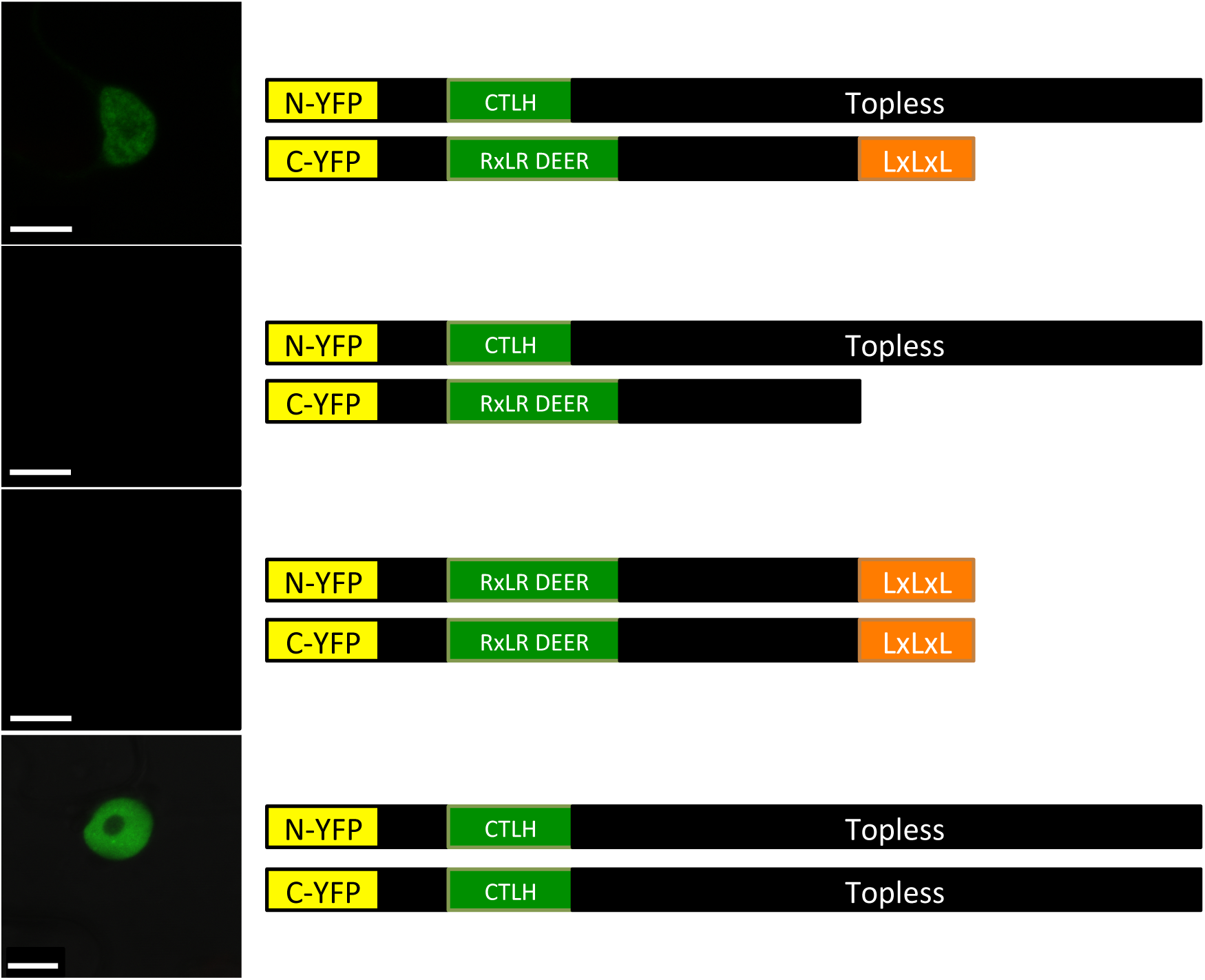
RxL21 interacts with TPL *in planta* by BiFC. The BiFC assay was conducted transiently in *N*. *benthamiana* epidermal cells by co-agroinfiltration of TPL(YFP^N^) and RxL21(YFP^C^), TPL (YFP^N^) and RxL21ΔEAR(YFP^C^), RxL21(YFP^N^) and RxL21(YFP^C^), TPL(YFP^N^) and TPL(YFP^C^). Infiltrated tissues were imaged at 48 hpi by confocal scanning laser microscopy for YFP fluorescence. The interaction between TPL and RxL21 was lost with deletion of the EAR motif. RxL21 does not appear to form dimers unlike TPL where strong fluorescence was observed in the nucleus. Scale bar = 10 μm.

**Figure S4.**
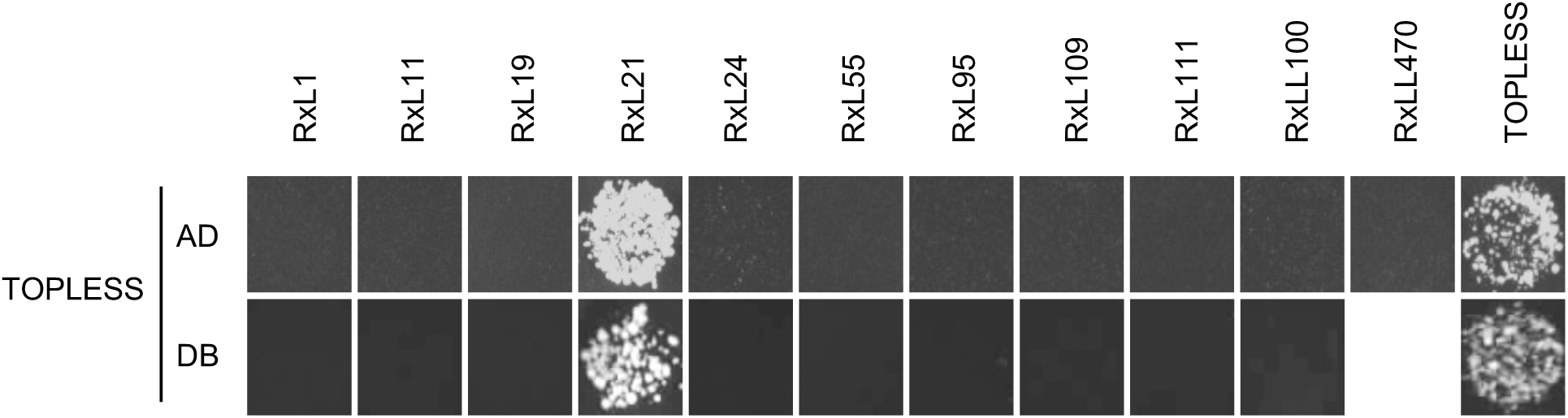
Interaction with TPL is specific to RxL21 amongst *Hpa* EAR-motif containing effectors. Y2H was performed using activation domain (AD)-TPL and DNA-binding domain (DB)-Effector constructs (top) and DB-TPL with AD-Effector constructs (bottom row). TPL interaction with HaRxL21 and TPL dimerisation were used as positive controls. Protein-protein interaction is shown by growth (indicating *GAL1::HIS3* reporter gene activation) on SC media lacking Leucine, Tryptophan and Histidine. All combinations tested also showed growth on –LW media indicating successful mating (not shown). The experiment was repeated on multiple plates with similar results.

**Figure S5.**
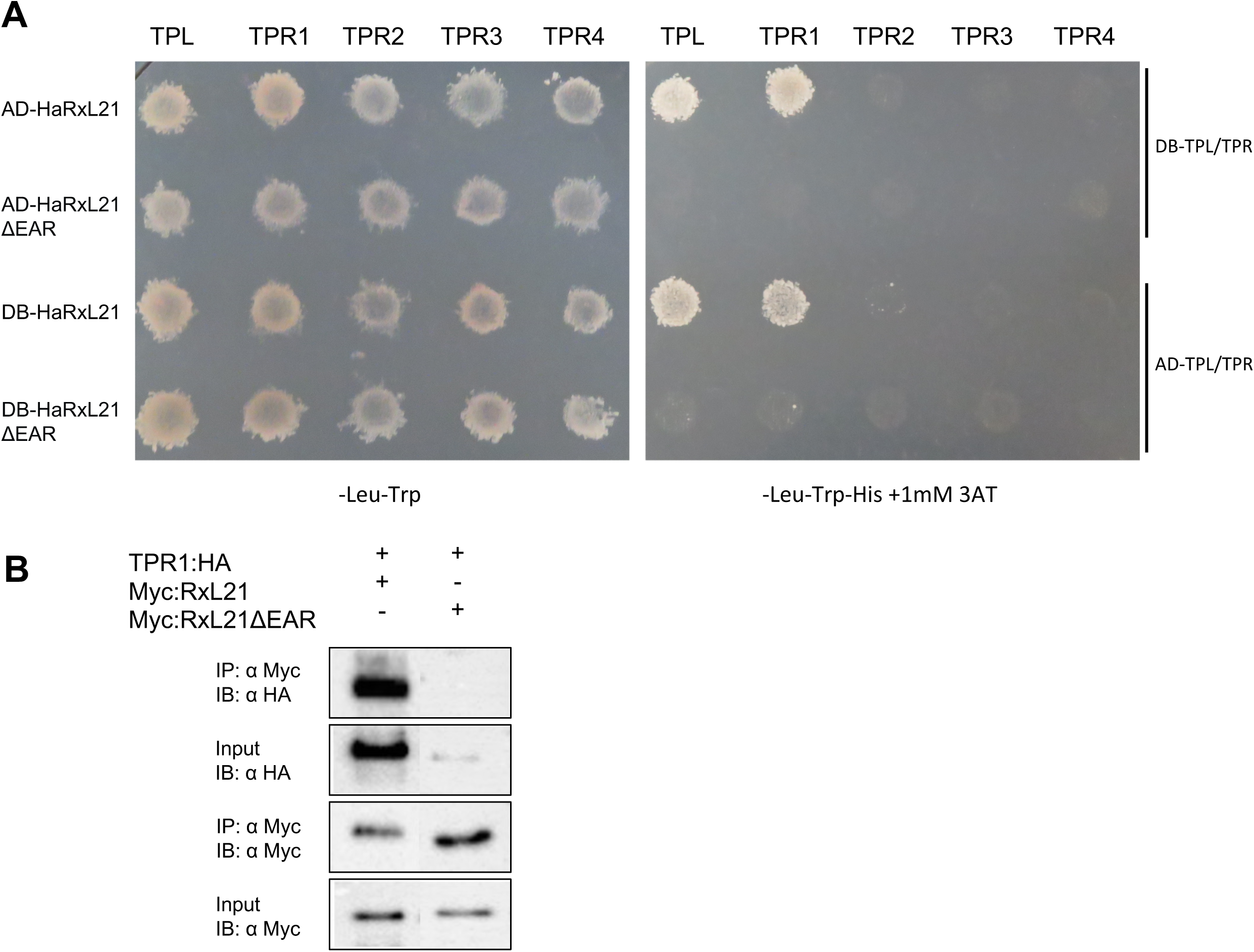
RxL21 interacts with TPR1. (A) RxL21 interacts with TPL and TPR1 by Y2H, indicated by growth on media lacking Leucine (Leu), Tryptophan (Trp) and Histidine (His). Growth on –Leu-Trp media indicates successful mating. Y2H was repeated in both directions, with both RxL21 and TPRs fused to AD; activation domain and DB; DNA binding domain. Y2H was repeated on multiple plates with similar results. (B) TPR1 interacts with RxL21 *in planta*. TPR1:HA with Myc:RxL21 or Myc:RxL21ΔEAR were transiently expressed in *N*. *benthamiana* leaves and harvested after 48 h. RxL21 expression was induced by 30μM β-estradiol 24 h prior to harvesting. α-myc beads were used for immunoprecipitations (IP). HA antibody was used to detect TPR1 immunoblots (IB) and α-myc antibody was used to detect RxL21 and RxL21ΔEAR.

**Figure S6.**
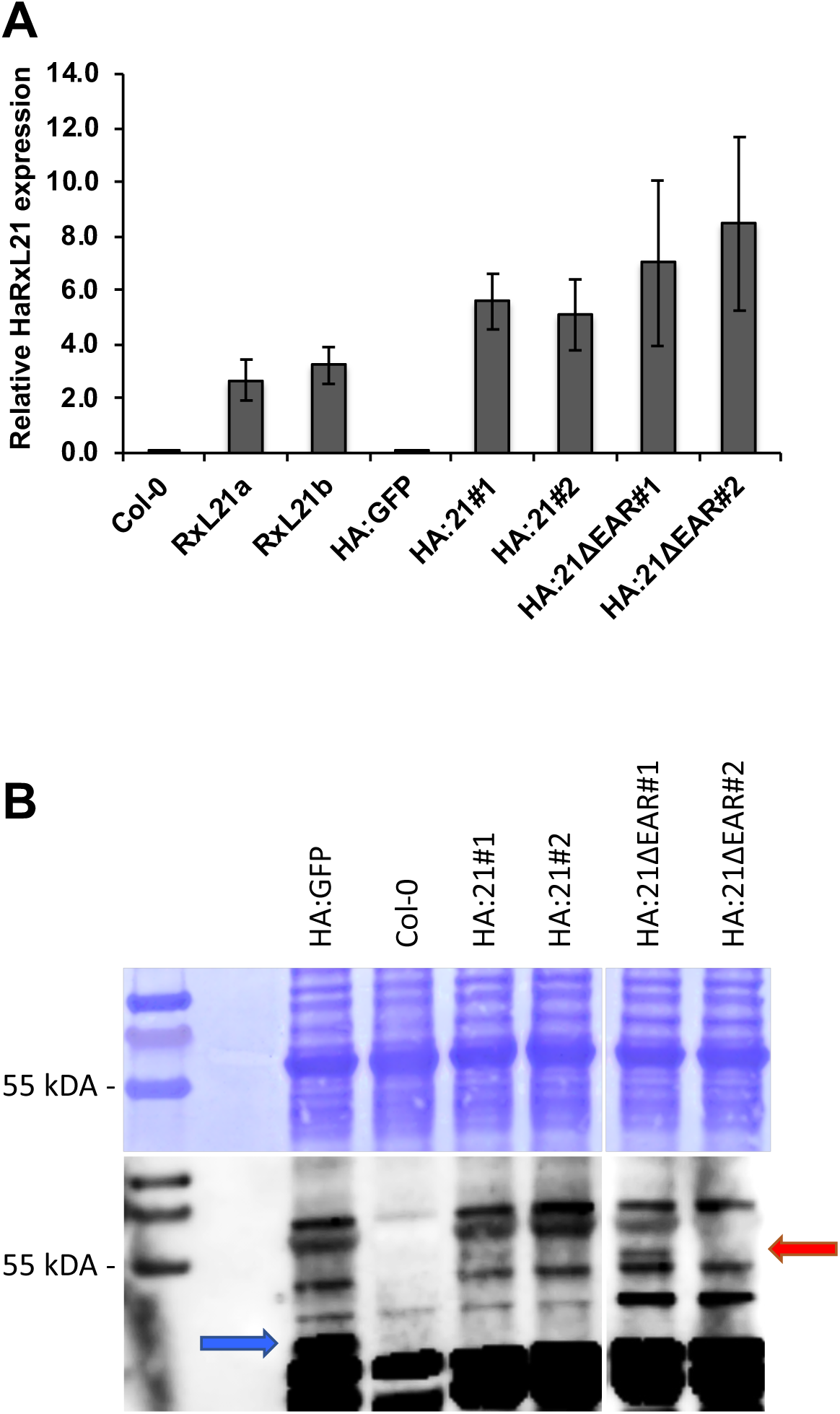
Gene and protein expression in 35S::HA::RxL21 lines. (A) Expression of *RxL21* by quantitative RT-PCR relative to *AtAct2* and *UBQ5*. Error bars show variance between technical replicates. Previously characterized 35S lines RxL21a/b were included for comparison. (B) Upper: Coomassie staining, lower: western blot using anti-HA. GFP and RxL21 bands are indicated by blue and red arrows respectively.

**Figure S7.**
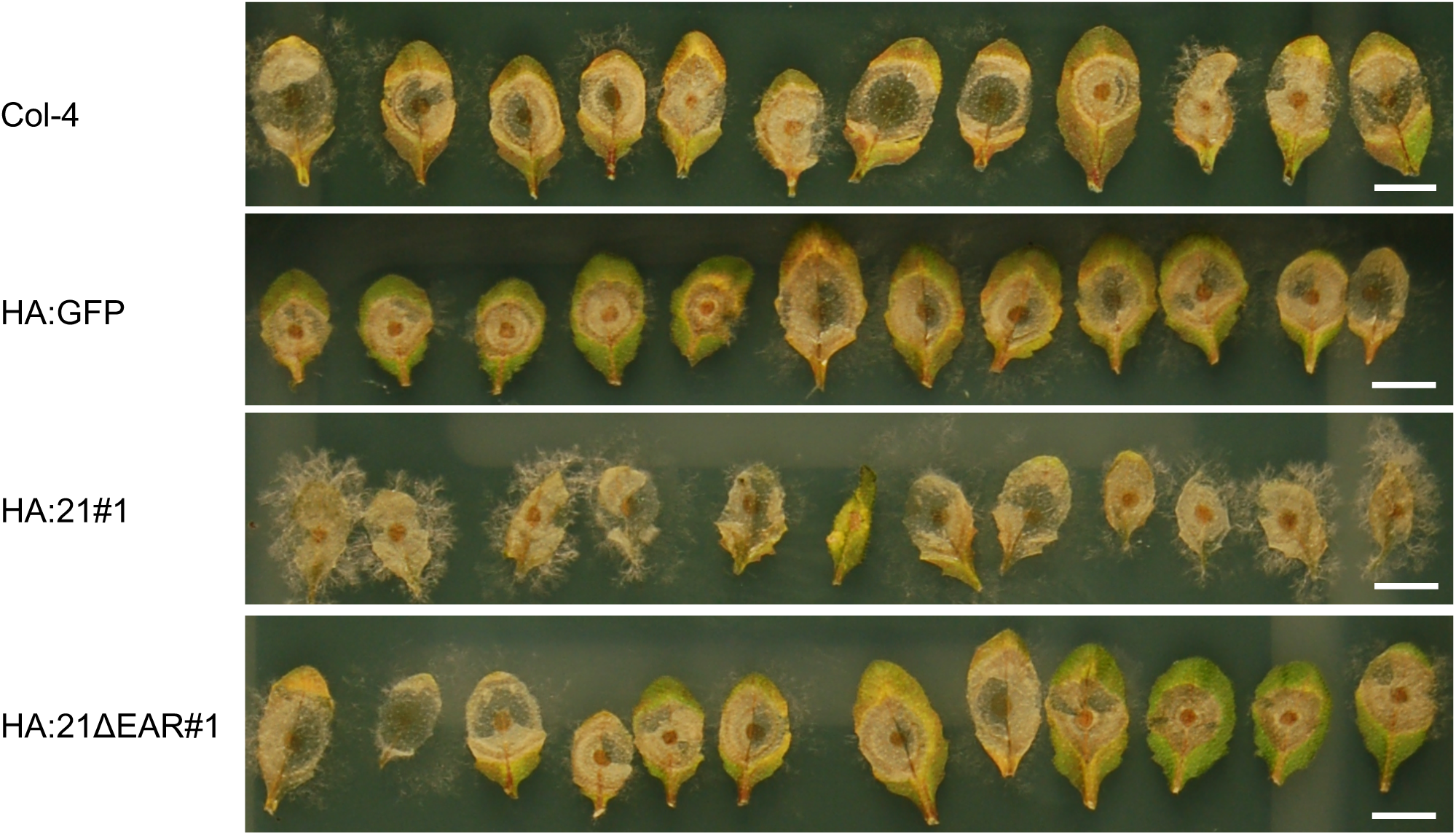
RxL21 lines show more visual sporulation after *Botrytis cinerea* infection. Photos taken 96 h post-infection with *Botrytis cinerea*. HA:21#1 appears to show more sporulation compared to Col-4, 35S::HA::GFP (HA:GFP) and HA:21ΔEAR#1. Scale bar is 1 cm.

**Figure S8.**
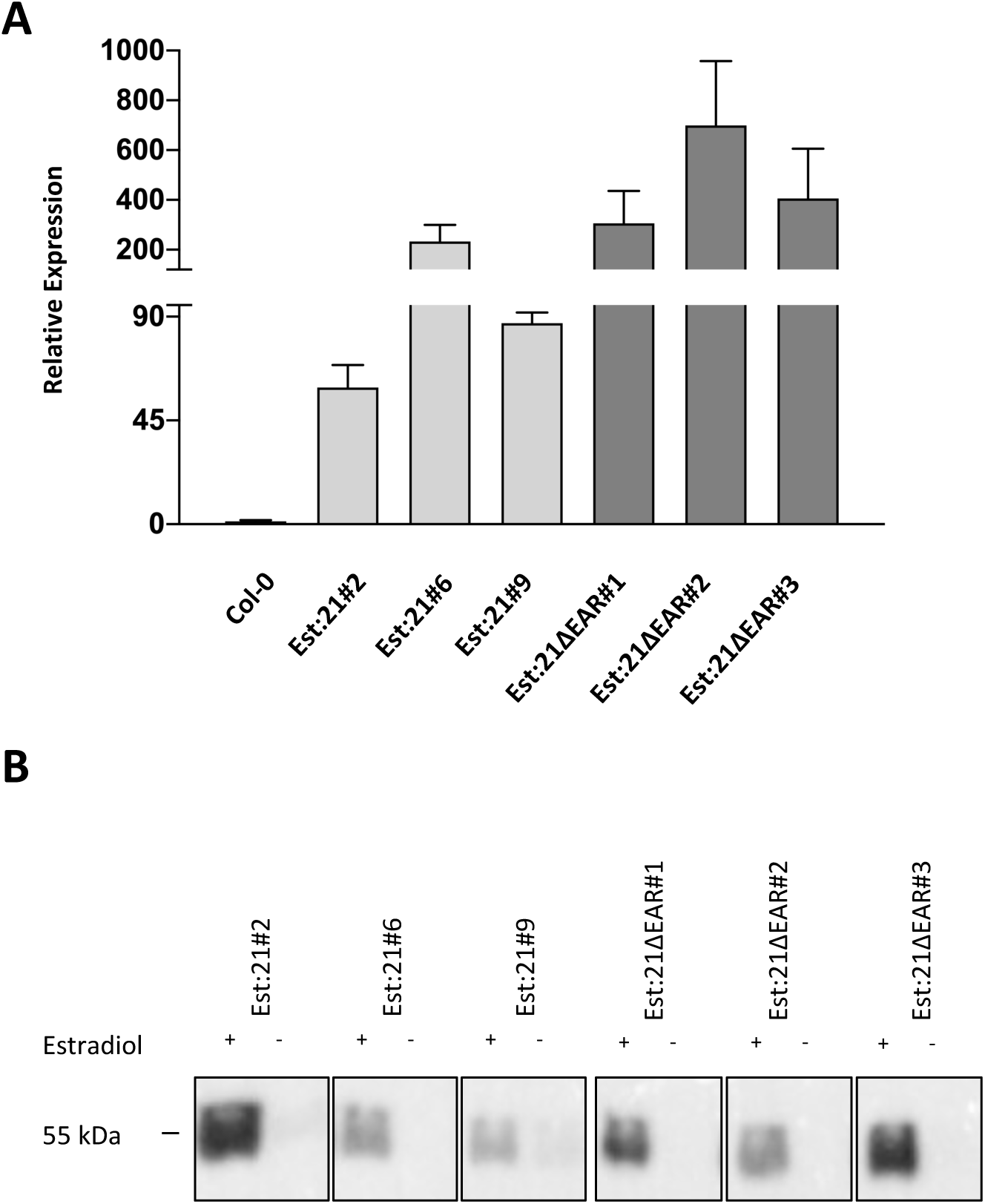
RxL21 expression in RxL21 and RxL21ΔEAR estradiol inducible lines. (A) Relative expression of *RxL21* in Arabidopsis plants expressing myc::RxL21 and myc::RxL21ΔEAR (under an estradiol (Est) inducible promoter) was determined by quantitative RT-PCR 18 h after induction with 30 μM estradiol. Expression levels were normalised to Arabidopsis *tubulin <u>4</u>*. Error bars show standard error between 3 biological replicates. (B) Anti Myc immunoblot showing Est-inducible expression of RxL21 and RxL21ΔEAR in Arabidopsis lines. Samples were taken 18 h after induction with 30 μM estradiol.

**Figure S9.**
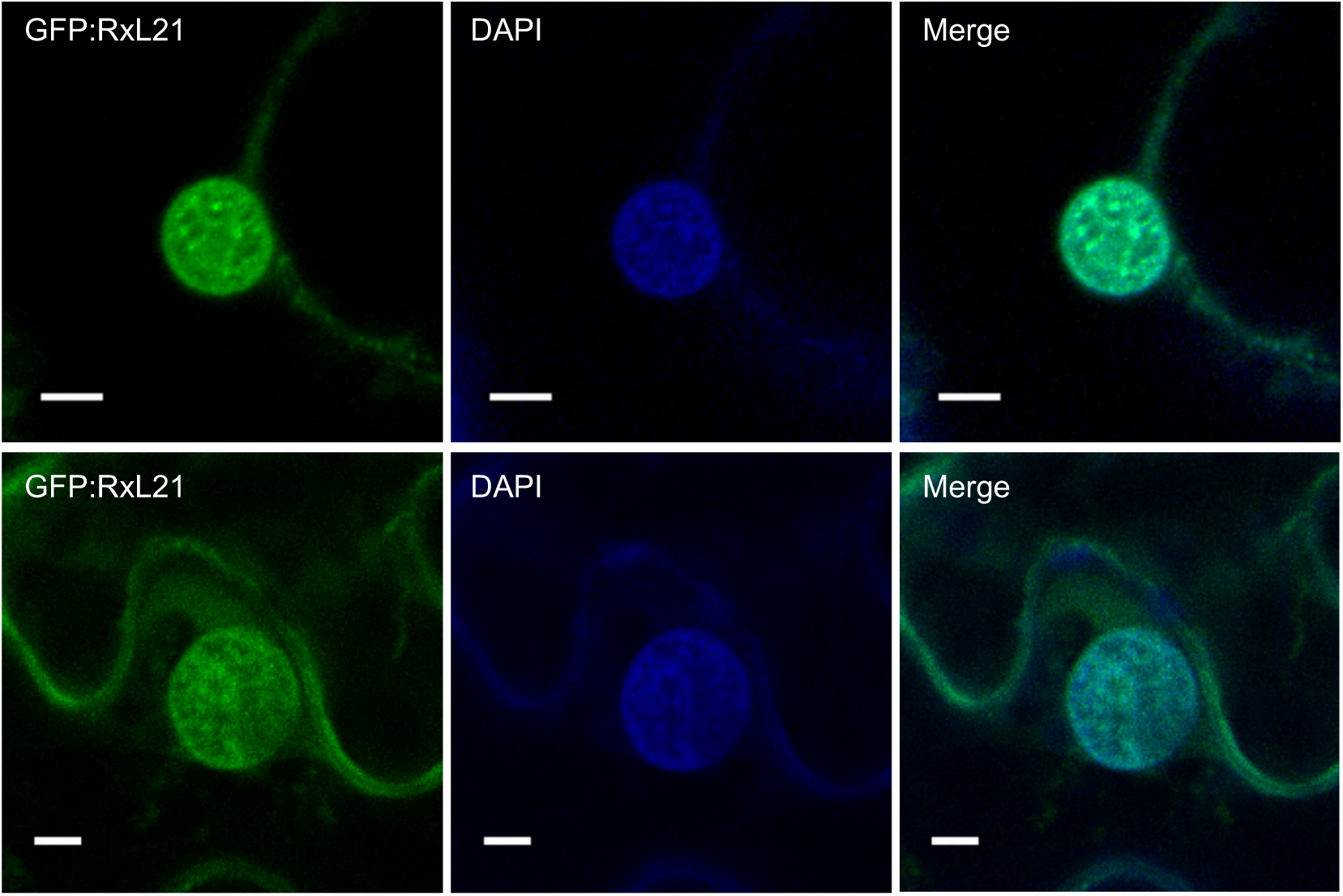
RxL21 co-localizes with DAPI in the nucleus. Transient expression of GFP:RxL21 using Agrobacterium mediated transformation of *N*. *benthamiana*. Immediately prior to imaging, leaves were stained by infiltration with DAPI (4′,6-diamidino-2-phenylindole). Scale bars are 5 μm.

**Figure S10.**
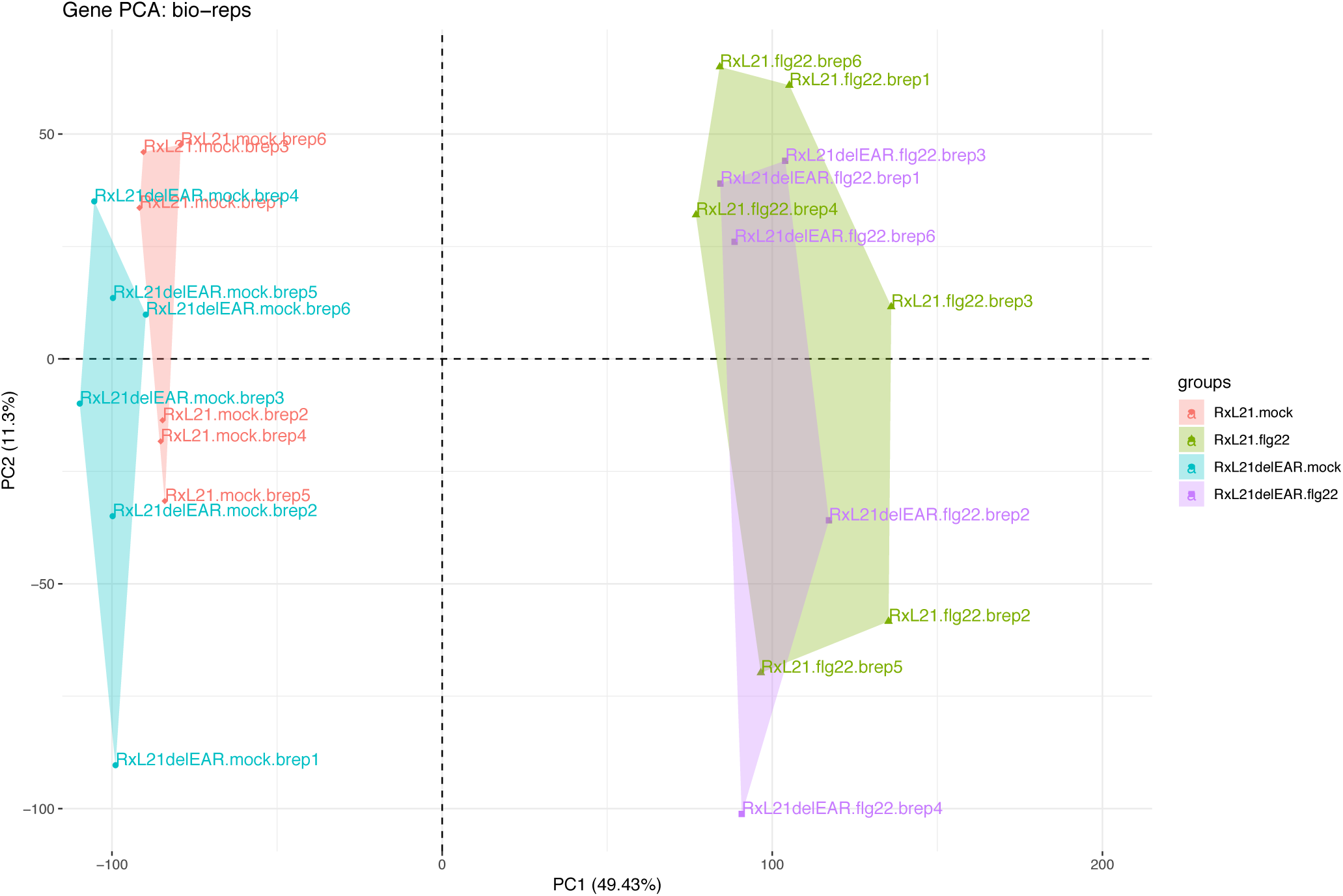
Principal component analysis showing variance between samples used for RNA-seq. A principal component analysis (PCA) plot characterizes the trends shown by the RNAseq data. Each point represents a sample (mean of 2 technical replicates) and colour characterizes the sample group. Groups consist of 6 bioreps (3 samples from each of two independent transgenic Arabidopsis lines expressing 35S:HA:RxL21 or 35S:HA:RxL21ΔEAR).

**Figure S11.**
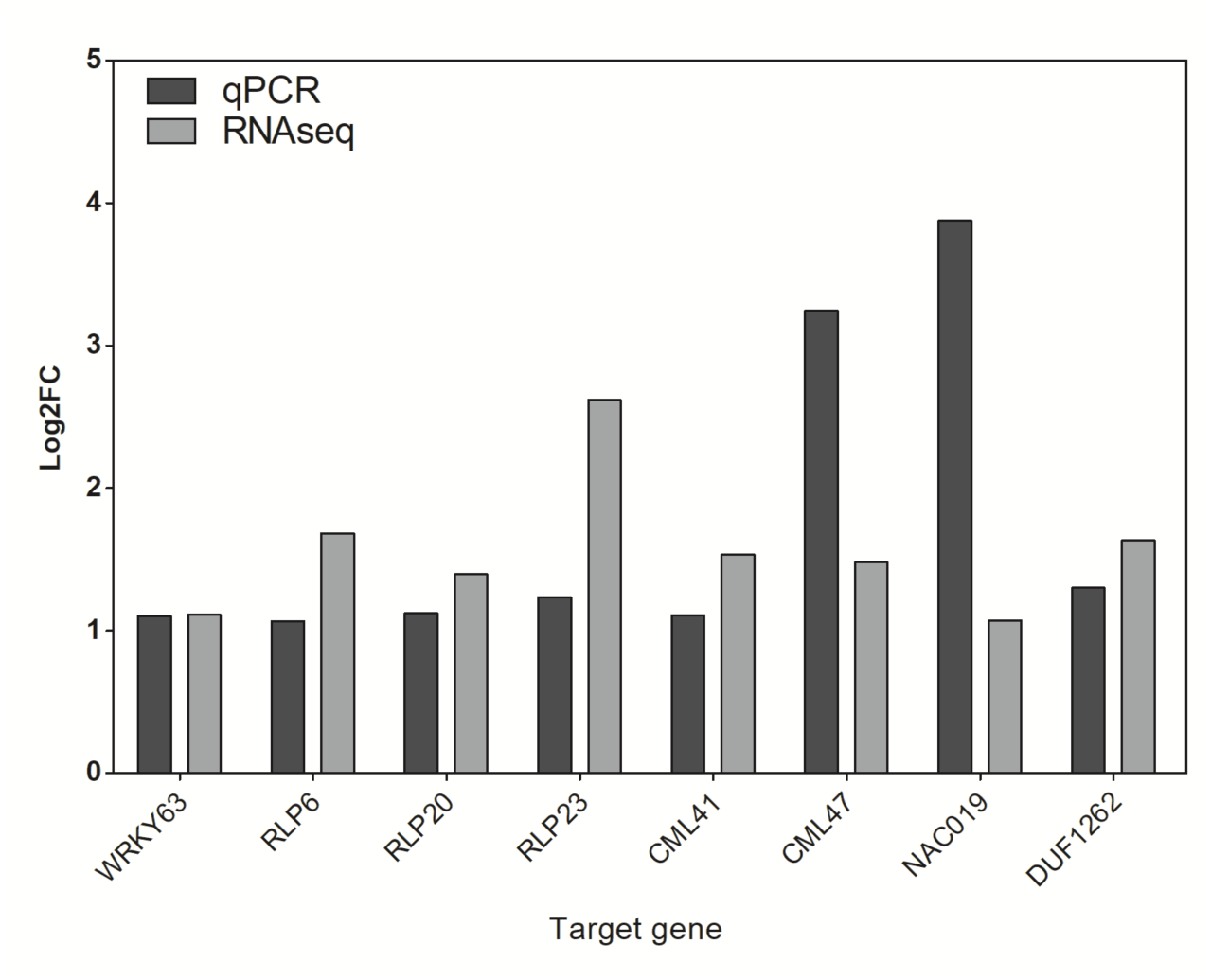
Comparison of RNA-seq expression data and qPCR data. Comparison of log_2_ fold change (log_2_FC) for the 8 selected genes between RxL21 and RxL21ΔEAR lines by qPCR compared to RNA-seq read counts. Mean fold change of three biological replicates is shown.

